# Improved gene annotation of the fungal wheat pathogen *Zymoseptoria tritici* based on combined Iso-Seq and RNA-Seq evidence

**DOI:** 10.1101/2023.04.26.537486

**Authors:** Nicolas Lapalu, Lucie Lamothe, Yohann Petit, Anne Genissel, Camille Delude, Alice Feurtey, Leen N. Abraham, Dan Smith, Robert King, Alison Renwick, Mélanie Appertet, Justine Sucher, Andrei S. Steindorff, Stephen B. Goodwin, Gert H.J. Kema, Igor V. Grigoriev, James Hane, Jason Rudd, Eva Stukenbrock, Daniel Croll, Gabriel Scalliet, Marc-Henri Lebrun

## Abstract

Despite large omics datasets, the establishment of a reliable gene annotation is still challenging for eukaryotic genomes. Here, we used the reference genome of the major fungal wheat pathogen *Zymoseptoria tritici* (isolate IPO323) as a case study to develop methods to improve eukaryotic gene prediction. Four previous IPO323 annotations identified 10,933 to 13,260 gene models, but only one third of these coding sequences (CDS) have identical structures. To resolve these discrepancies and improve gene models, we generated full-length transcripts using long-read sequencing. This dataset was used together with other evidence (RNA-Seq transcripts and protein sequences) to generate novel *ab initio* gene models. The selection of the best structure among novel and existing gene models was performed according to transcript and protein evidence using InGenAnnot, a novel bioinformatics suite. Overall, 13,414 re-annotated gene models (RGMs) were predicted, including 671 new genes among which 53 encoded effector candidates. This process corrected many of the errors (15%) observed in previous gene models (coding sequence fusions, false introns, missing exons). While fungal genomes have poor annotations of untranslated regions (UTRs), our Iso-Seq long-read sequences outlined 5’ and 3’UTRs for 73% of the RGMs. Alternative transcripts were identified for 13% of RGMs, mostly due to intron retention (75%), likely corresponding to unprocessed pre-mRNAs. A total of 353 genes displayed alternative transcripts with combinations of previously predicted or novel exons. Long non-coding transcripts (lncRNAs) and double-stranded RNAs from two fungal viruses were also identified. Most lncRNAs corresponded to antisense transcripts of genes (52%). lncRNAs that were up or down regulated during infection were enriched in antisense transcripts (70%), suggesting their involvement in the control of gene expression. Our results showed that combining different *ab initio* gene predictions and evidence-driven curation using InGenAnnot improved the quality of gene annotations of a compact eukaryotic genome. Our analysis also provided new insights into the transcriptional landscape of *Z. tritici*, helping develop an increasingly complex picture of its biology.

## Introduction

Predicting genes in eukaryotic genomes is a challenging process [1], particularly for fungi with compact genomes. The quality of a genome annotation depends on supporting evidence for coding regions, splice junctions and on the algorithms used to derive patterns for predictions [2]. Several drawbacks for gene annotation were identified in eukaryotic genomes such as the complexity of their gene structure, with introns difficult to predict without experimental transcript evidence, as well as the quality of genome assembly when fragmented in contigs. In fungi, genes are generally close to each other, and frequent overlaps between adjacent transcripts have been observed [3]–[5]. In addition, fungi have shorter introns (averaging 70-100 bp depending on the species, [6]) compared to other eukaryotes. These particularities of fungal genomes require specific training of *ab-initio* prediction software and development of fungal-specific pipelines [7]–[15]. Long-read sequencing is now used to provide full genome assemblies, reducing drawbacks due to genome fragmentation into contigs. Experimental transcript evidence has also been improved using transcripts assembled from RNA-seq short reads, providing large transcript datasets for gene annotation/curation. Iso-Seq long-read sequencing now provides full-length transcript sequences that bypass problems observed with the assembly of RNA-Seq short reads such as chimeric transcripts covering adjacent genes [16]. Iso-Seq also provides transcript isoforms allowing the identification of alternative start, stop and splicing events. Nevertheless, RNA-Seq reads are still required to quantify the relative abundance of Iso-Seq transcript isoforms, since Iso-Seq is not quantitative and could reveal rare transcripts likely resulting from errors of the transcriptional machinery [17]. Combining these two types of transcript sequencing is needed to avoid drawbacks from each technique [18]. Other omics methods such as transcription start site sequencing (TSS-seq) or cap-analysis gene expression sequencing (CAGE-seq) are now available for precise definition of transcript start sites, but these applications are still limited to model organisms [19], [20].

We have chosen the reference genome of the major fungal wheat pathogen *Zymoseptoria tritici* (isolate IPO323) as a case study to improve methods for eukaryotic gene prediction and curation. *Z. tritici* is an ascomycete (class Dothideomycetes, [21]) that causes a major foliar disease of bread and durum wheat (*Septoria tritici* blotch [22]). The first *Z. tritici* genome sequence was obtained in 2011 for the bread wheat-infecting European reference isolate IPO323 using Sanger sequencing [23]. This complete genome sequence from telomere to telomere has a size of 39.7 megabases (Mb) and is composed of 13 core chromosomes (CCs) and 8 accessory chromosomes (ACs). Chromosome-scale genome assemblies of 22 additional *Z. tritici* isolates from different geographic origins were obtained using long-read sequencing [24], [25], [26], as well as the genome sequences of four related species of *Zymoseptoria* (*Z. ardibilae, Z. brevis, Z. passerinii, Z. pseudotritici*) [25]. A large proportion of the IPO323 *Z. tritici* genome is composed of transposable elements (TEs, 17% to 20%, [27][28]), while the TE content of other isolates varied between 14% and 21.5% [24], [29], [30].

Currently, four annotations of the IPO323 *Z. tritici* genome are available. The first was generated by the Joint Genome Institute in 2011 (JGI, [23]). The second annotation was performed at the Max Planck Institute for Evolutionary Biology in 2015 (MPI, Germany, [28]). Two other annotations were generated in 2015 at Rothamsted Research Experimental Station (RRES,[31]) and the Centre for Crop & Disease Management of Curtin University. Large discrepancies were observed across annotations, both in gene numbers (10,933 to 13,260) and gene structures (30% of coding sequences (CDS) with identical structures). In addition, some genes that are important for the infection process of *Z. tritici* were not predicted. For example, the effector-encoding gene *Avr-Stb6* was located near the telomere of chromosome 5 by quantitative trait locus (QTL) mapping and genome-wide association study (GWAS), but it was not predicted in existing IPO323 annotations [32]. Indeed, it was identified by translating all possible ORFs from the region, and its overall structure (start, stop, two introns) was only predicted using infection-related RNA-seq data. Clearly, the complete coding potential of this genome still has not been identified despite the four thorough annotations that have been developed over the past dozen years.

To address this problem, we established a novel strategy to annotate a compact eukaryotic genome using *Z. tritici* as a case study. For this process we generated a large set of full-length cDNA sequences using PacBio Iso-Seq long reads [33], [34]. We also developed a novel suite of tools, InGenAnnot, to compare genes models predicted by different *ab initio* software and to select the best gene model according to transcript (RNA-Seq, Iso-Seq) and protein evidence. A novel set of 13,414 improved gene models was generated. Comparing this annotation to other annotations revealed systematic errors in previous gene models. Full-length cDNA sequences were also used to identify alternative transcripts and long, non-coding RNA (lncRNA), improving our understanding of the transcriptional landscape of *Z. tritici*.

## Materials and Method

### Available *Z. tritici* IPO323 gene annotations

Currently, four annotations of the *Z. tritici* IPO323 genome are available. The first, with 10,933 gene models, was developed in 2011 by the Joint Genome Institute with *ab initio* tools FGENESH and Genewise [8] using EST (expressed sequence tag) and proteome evidence (JGI, [23]). The second annotation was performed in 2015 by the Max Planck Institute, resulting in 11,839 gene models (MPI, Germany, [28]) identified with the Fungal Genome Annotation pipeline [35]. This pipeline uses *ab initio* tools GeneMark-ES, GeneMark-HMM [13] and Augustus [12] combined by EVidenceModeler [36] with RNA-Seq evidence and keeping as much as possible of the first annotation provided by JGI. The third annotation was generated in 2015 by the Rothamsted Research Experimental Station (UK) with 13,862 gene models (RRES, [31]) obtained with the *ab initio* tool MAKER-HMM [11] and RNA-Seq evidence. The fourth annotation published in 2015 by the Centre for Crop & Disease Management, Curtin University. (CURTIN, Australia) with 13,260 gene models, was obtained with *ab initio* tool CodingQuarry [37] and RNA-Seq evidence. All gene files used in the annotations by JGI, MPI, RRES and CURTIN have been made easily accessible (https://doi.org/10.57745/CVIRIB) and can be displayed with a dedicated genome browser (https://bioinfo.bioger.inrae.fr/portal/genome-portal/12) or on the new IPO323 genome web portal at JGI (https://mycocosm.jgi.doe.gov/Zymtr1/Zymtr1.home.html).

### Fungal Isolate, RNA extraction, PacBio Iso-Seq and Illumina RNA-Seq libraries

The reference isolate of *Z. tritici* IPO323 [23] was stored at -80°C as a yeast-like cell suspension (10^7^ cells/mL in 30% glycerol). *Z. tritici* was grown at 18°C in the dark on solid (Yeast extract Peptone Dextrose (YPD) agar) or liquid (Potato Dextrose Broth (PDB)) media. For RNA production, *Z. tritici* isolate IPO323 (4-day-old yeast-like cells diluted to 10^5^ cells/mL final) was cultivated in 75-mL agitated liquid cultures (500 mL Erlen flasks, 150 rpm) at 18°C in the dark for 4 days. Different media were used (Table S3) including Glucose-NO_3_ synthetic medium defined as MM-Zt [38]. MM-Zt was modified by replacing glucose (10 g/L) by different carbon sources (Xylose, Mannitol, Galactose, Sucrose at 10 g/L)). Histone Deacetylase inhibitors such as trichostatin ((TSA, Sigma T8552, 1 μM final) and SAHA (SAHA, Sigma SML0061, 1 mM final) were added to MM-Zt to express genes located in genomic regions with repressive chromatin marks [39]. The composition of complex media (Yeast-Peptone-Dextrose: YPD, Potato-Dextrose-Both: PDB, Glycerol-Nitrate: AE) was already described [40]. Cultures of IPO323 in YPD and PDB were performed at 18°C and 25°C, while AE cultures were performed only at 18°C. A total of 14 culture conditions was used for RNA production (Table S3). All cultures for RNA-Seq were performed in triplicate. Cultures were centrifuged at 3000 rpm for 10 minutes and mycelium pellets were washed with water and frozen with liquid nitrogen. Frozen mycelium was lyophilized and kept at -80°C until extraction. RNAs were extracted using the Qiagen Plant RNeasy Kit according to the manufacturer’s protocol (Ref. 74904, Qiagen France SAS, Courtaboeuf, France). Preparation and sequencing of PacBio Iso-Seq libraries were performed by the INRAE platform Gentyane (http://gentyane.clermont.inrae.fr). The SMARTer PCR cDNA Synthesis Kit (ref 634926, Clontech, Mountain View, CA, USA) was used for polyA-primed first-strand cDNA synthesis followed by optimized PCR amplification and library preparation using the SMRTbell Template Prep Kit (ref 101-357-000, Pacific Bioscience, Menlo Park, CA, USA) according to manufacturer protocols. The cDNA libraries were prepared without size selection and bar coded for multiplexing. Sequencing was performed on a PacBio SEQUEL (version 1). Illumina RNA-seq single-stranded libraries were prepared using the NEBNext Poly(A) mRNA Magnetic Isolation Module (NEB #E7490, New England BioLabs, Ipswich, Massachusetts, USA) and the NEBNext Ultra II Directional RNA Library Prep Kit for Illumina (<u>NEB #E7765</u>, New England BioLabs, Ipswich, Massachusetts, USA). Custom 8-bp barcodes were added to each library during the preparation process. Pooled samples were cleaned with magnetic beads included in the library preparation kit. Each pool was run on a lane of Illumina HiSeqX (Illumina, San Diego, California, USA) using a 150-cycle paired-end run

### Processing of RNA-seq sequences

RNA-Seq data were cleaned and trimmed with Trimmomatic (v 0.36) [41]. The cleaned sequences were then mapped to the *Z*.*tritici* IPO323 genome using STAR (v 2.5.1b, --alignIntronMin 4 -- alignIntronMax 5000 -- alignMatesGapMax 5000) [42]. Wig files of uniquely mapped reads were converted to BigWig files with wigToBigWig (v4). StringTie (v2.1.1) [43] was then used to assemble the mapped RNA-Seq reads into transcripts with different parameters depending on the depth of sequencing of libraries and their type (-m 150 --rf --g 0 -f 0.1 -a 10 -j 2 or -j 4). The Trinity script inchworm_transcript_splitter.pl (version 2.8.5) [44] was used to split the transcripts with non-uniform coverage based on the Jaccard clip method. Clipped transcripts were extracted with home-made scripts and clustered with Stringtie and associated bam files to obtain transcripts per million (TPM) counts. All libraries were concatenated into one gff file without merge to avoid loss of information by fusion of small transcripts into larger ones due to the large number of genes in the *Z. tritici* genome with overlapping untranslated regions (UTRs).

### Processing of Iso-Seq sequences

Iso-Seq raw data were processed with the Iso-Seq V3.2 pipeline from PacBio generating polished Circular Consensus Sequences (CCS). CCS were then mapped to the *Z. tritici* IPO323 genome with Gmap (2019-01-31) [45] and unmapped, low-mapping-quality (≤0) or multi-mapped CCS were filtered out. The CupCake package (v10.0.0, https://github.com/Magdoll/cDNA_Cupcake) filtered the isoforms, removing the less-expressed and degraded transcripts using the following tools: *collapse_isoforms_by_sam*.*py, get_abundance_post_collapse*.*py, filter_by_count*.*py, filter_away_subset*.*py*. Readthrough transcripts were removed using the previous annotations (MPI, JGI, CURTIN, RRES) with BEDTools intersect [46] with an an overlap of 100% for full coding sequences (CDS) (-F 1.0) and the same strand (-s)) of at least 2 CDS. Transcripts mapped on the mitochondrial genome were filtered out as well. Subsequently, all libraries were processed with *chain_samples*.*py* from CupCake and clustered for stringent selection. Splicing junctions obtained by STAR (SJ.out.tab files) from Illumina RNA-Seq libraries were used to filter out isoform transcripts with unsupported junctions. Finally, long-read transcripts fully spanning transposable elements were removed with BEDTools, giving the final set of transcript evidence.

### Gene prediction and selection of the best gene models

Two gene predictors, Eugene v1.6.1 [10], and LoReAn v2.0 [47], handling long-read transcript sequences as evidence, were used to perform new annotations. Eugene was launched with the provided fungal parameters (WAM fungi matrix) and trained with a dataset of proteins from four genomes of species phylogenetically related to *Z. tritici*: *Cercospora beticola* (GCF_002742065.1_CB0940_V2); *Ramullaria collo-cygni* (GCF_900074925.1_version_1); *Zasmidium cellare* (GCF_010093935.1_Zasce1); and *Sphaerulina musiva* (GCF_000320565.1_Septoria_musiva_SO2202_v1.0). Gene structures were predicted with assembled transcripts from RNA-Seq and a dataset of Dothideomycetes proteins obtained from Uniprot without *Zymoseptoria* sequences to avoid inference with gene models to be improved. Filtered Iso-Seq transcripts were used as strongly weighted evidence in model prediction with the parameter “`est_priority=2`”. LoReAn was launched in fungus mode with the Augustus retraining mode using the same Dothideomycetes Uniprot dataset without *Zymoseptoria* sequences and the same Iso-Seq transcript dataset used for Eugene. RNA-Seq data were used as a merged mapping file (BAM) by the pipeline to assemble transcripts and detect splicing sites. The new and previous gene datasets cleaned for TEs with *ingenannot filter* were annotated for annotation edit distance (AED) [48] scores using *ingenannot aed* with a fungal protein dataset without any *Zymoseptoria* species, selected Iso-Seq and RNA-Seq transcripts. AED were computed on gene models only with “`--aed_tr_cds_only”` to avoid bias between datasets with or without UTR annotations and with “`--penalty_overflow 0.25`” to penalize gene models with splicing junctions that lacked support evidence. The best gene models were selected with *ingenannot select* based on a AED of ≤0.3 for transcript or an AED of ≤0.1 for protein evidence. Gene models failing the AED threshold, but contained in clusters with at least 4 predictions from independent annotations were retained, but partial gene models (no ATG nor stop codon) were removed. The high number of annotation sources (6) and selection of loci detected by 4 independent annotations, allow us to use stringent AED thresholds, limiting selection of annotation-specific gene models to well supported structures.

Potential new gene effectors predicted with *ingenannot rescue_effector*, were added to the final set. Transcripts not co-localizing with a selected gene model were tested in 3 frames to analyse the predicted peptides with the same criteria used to detect small, secreted proteins (SSP) as described below. UTRs were inferred in two passes with the *ingenannot utr_refine*. First, after deleting all previously annotated UTRs and inferring new coordinates from a filtered set of Iso-Seq transcripts. Second, by inferring UTRs with a filtered set of RNA-seq assembled transcripts, considering only transcripts with no UTRs from the first step. Both sets were established with *the ingenannot isoform_ranking* for filtering and ranking UTR isoforms based on RNA-Seq evidence.

Gene models from each annotation were compared using their AED scores with *ingenannot aed_compare* and specific/shared gene models were identified using *ingenannot compare*. BUSCO [49] analyses with ascomycota_odb10 were performed to evaluate the completeness of datasets.

### Functional annotation and prediction of secreted proteins

Functional annotations of genes obtained using Interproscan 5.0 [50] and Blastp [51] (e-value <1e-5) against the NCBI nr databank were then used to perform Gene Ontology annotation [52] with Blast2GO [53]. Secretomes and effectors were annotated as described in [54]. The secretome was predicted by a combination of TMHMM (v.2.0) [55], SignalP (v4.1) [56] and TargetP (v1.1b) [57] results with the following criteria: no more than one transmembrane domain and either a signal peptide or an extracellular localization prediction. The SSP repertoire was predicted by applying a size cut-off of 300 amino acids to the predicted secretome and keeping only proteins predicted as effectors by EffectorP (v2.0).

### Analysis of Iso-Seq transcript isoforms

The annotation of transcript isoforms was performed with sqanti3 [58] using Iso-Seq transcripts, previously established to infer UTRs, filtered for UTR length isoforms and low expression levels (less than 10% of total RNA-Seq reads), using the *ingenannot isoform_ranking* tool. RNA-Seq reads were mapped to Iso-Seq transcripts with RSEM v1.3.3 [59] and Differential Isoform Usage (DIU) performed with tappAS [60] with annotations obtained from sqanti3.

### Detection of antisense and lncRNA Iso-Seq transcripts

Iso-Seq transcripts annotated as antisense and intergenic with sqanti3 were selected as Putative long non-coding (lnc) RNAs. Then transcripts shorter than 1 Kb in length [61], overlapping with TEs and containing an open reading frame (ORF) longer than 100 amino acids predicted with getorf by EMBOSS [62] were discarded. The resulting “non-coding” transcripts were annotated with CPC2 [63], and only transcripts without an ORF with a PFAM domain were kept as lncRNAs. featureCounts (v1.5.1) [64] was used to count reads per transcript, followed by differential expression analysis by edgeR [65] with the SARTools package (v1.6) [66].

### Detection of polycistronic Iso-Seq transcripts

For detecting polycistronic mRNAs,, read-through Iso-Seq transcripts that were previously filtered out were merged to obtain the global counts of genes that were potentially co-transcribed. To establish a robust list of co-transcribed multi-gene loci, readthrough transcripts were filtered with the gene reannotation dataset and their Iso-Seq transcripts used as evidence. Only polycistronic mRNAs supported by independent long-read single transcripts for each gene were conserved and considered as reliable. Detection of overlaps between transcripts and annotations was performed with intersect using BEDTools [46].

### Identification and annotation of mycoviruses

Iso-Seq transcripts not mapping to the *Z. tritici* IPO323 reference genome were clustered with blastclust. Similarities with known sequences were analysed by *blastn* search against the NCBI nr database. Reconstruction of the full-length sequences of viruses was performed by de-novo assembly with SPAdes (v3.15.4) [67]. RNA-dependent RNA polymerase sequences from narnaviruses related to Zt-NV1 were retrieved from NCBI and analyzed using Phylogeny.fr [68]. Alignment of protein sequences was performed with Muscle 3.8.31 and curated by G-blocks. The phylogenetic analysis was performed using PhyML 3.1 and the phylogenetic tree was drawn with TreeDyn 198.3.

## Results

### Comparison of existing *Z. tritici* IPO323 genome annotations

The four *Z. tritici* IP0323 genome annotations (MPI, JGI, RRES, and CURTIN), filtered out for TE-encoding genes, were clustered into 13,225 metagenes corresponding to 26,224 distinct gene models using only their CDS as reference. Metagenes of InGenAnnot are clusters of overlapping genes transcribed from the same strand and corresponding to the “gene locus” defined in ParsEval [69]. To compare the structure of gene models from different annotations, we defined three categories: a) identical gene models (exactly the same CDS); b) dissimilar gene models (same metagene but different CDS); and c) specific gene models (CDS found only by one annotation at a given locus). Only 3,618 identical gene models were shared along the four annotations. When omitting the JGI annotation, the number of identical gene models among the MPI, RRES, and CURTIN annotations increased to 6,816 (Figure 1a). The highest numbers of identical gene models between two annotations were observed for MPI-RRES (8,442), RRES-CURTIN (8,289), and MPI-Curtin (7,981), while the lowest numbers of identical gene models were observed between JGI and the three other annotations (4,495, 4,621 and 5,276 for JGI-Curtin, JGI-MPI and JGI-RRES respectively). The RRES and CURTIN annotations displayed the highest numbers of specific gene models (593 and 436, respectively), while the MPI annotation displayed the lowest number of specific gene models (12). The JGI and CURTIN annotations displayed a higher number of dissimilar gene models (4,752 and 3,844, respectively) compared to the other annotations (2,367 and 1,871 for RRES and MPI, respectively; Figure 1).

**Figure 1.**
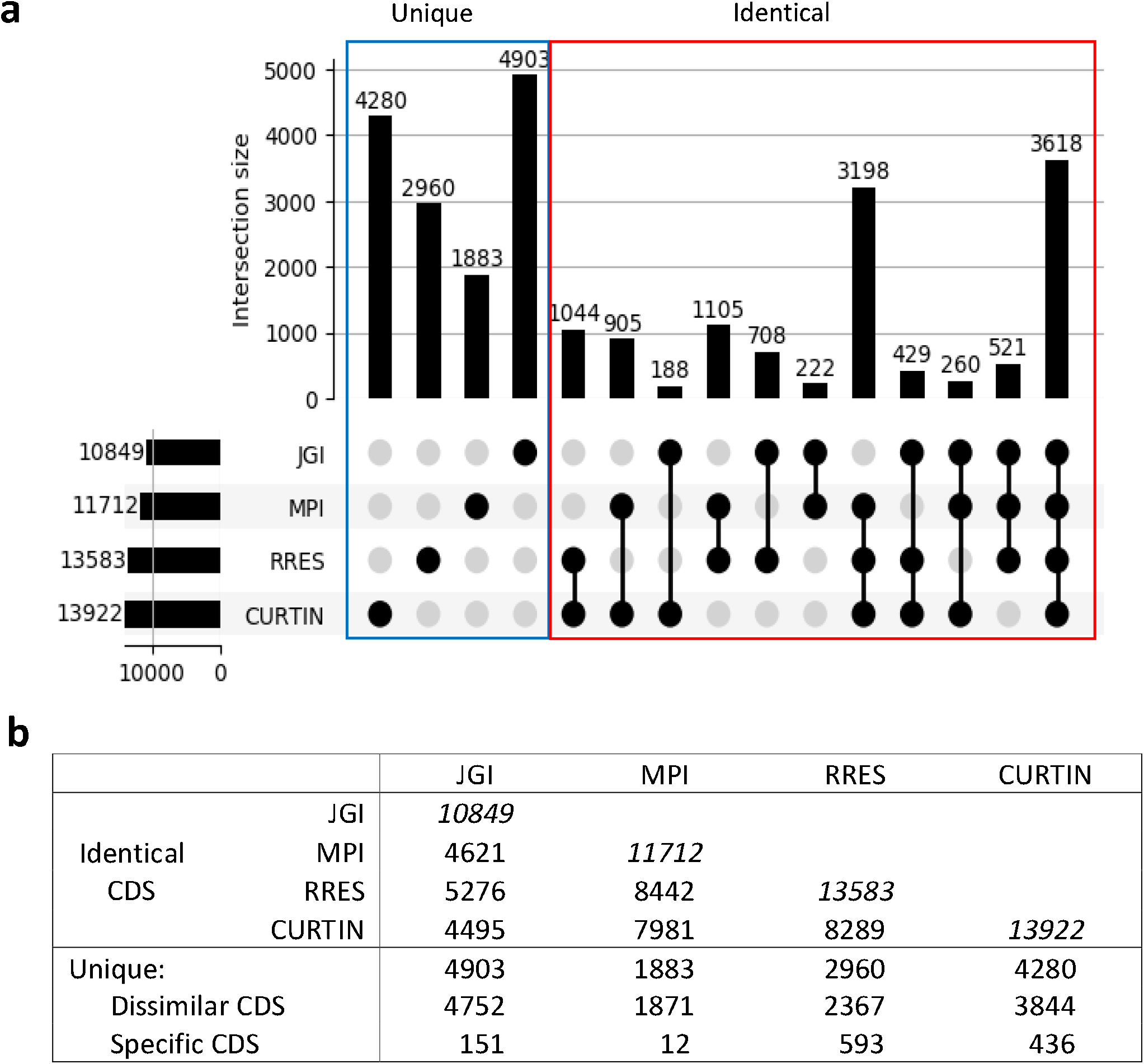
Comparison of *Zymoseptoria tritici* reference isolate IPO323 genome annotations. a) Upset plot of the gene models from the four annotations of IPO323 (JGI, MPI, RRES and CURTIN). Number of gene models with identical coding sequences (CDS). b) Comparison of IPO323 gene annotations. Number of CDS in each annotation. Identical CDS: identical CDS at a given locus. Unique Dissimilar CDS: at a given locus, a CDS is predicted by at least one other annotation, but they differ in their structure. Unique Specific CDS: at a given locus, a single CDS is predicted by a single annotation.

Despite the low numbers of identical gene models across annotations, basic genomic statistics were similar (Table S1). Still, the number of mono-exonic gene models was higher (1.4 to 1.8 fold) in the RRES and CURTIN annotations compared to those by the JGI and MPI. Most of these mono-exonic gene models were only predicted *ab initio* (without transcript or protein evidence) and they were often specific to a given annotation. The average size of gene models also differed between MPI and the other annotations (1465 bp compared to 1300 bp). We suspected that this difference could result from longer gene models corresponding to the fusion of two or more distinct adjacent gene models that were predicted as single genes by other annotations. Indeed, 533 and 801 gene fusions were detected in the MPI annotation, corresponding to at least two distinct adjacent gene models in the RRES and CURTIN annotations, respectively.

The chromosomal localization of gene models was compared across the four annotations (Table S2). The JGI, MPI and CURTIN gene models exhibited a similar distribution across chromosomes, while the RRES annotation displayed twice as many gene models on accessory chromosomes compared to other annotations. Overall, the low number of identical gene models across annotations (27% of metagenes) likely resulted from drawbacks of each annotation pipeline. For example, we identified many gene fusions in the MPI and JGI annotations. We also detected annotation-specific mono-exonic genes in the CURTIN and RRES annotations. These drawbacks resulted in the accumulation of both wrong and specific gene models in each annotation.

To circumvent these problems, we generated a novel annotation of the IPO323 genome relying on broad transcriptional evidence. This strategy required the construction of an expression dataset using both publicly available single-stranded RNA-Seq datasets, including wheat leaf infection kinetics, and newly generated datasets using both long-read sequencing (PacBio Iso-Seq: Iso-Seq) and short-read sequencing (single-stranded Illumina RNA-Seq: RNA-Seq) (Table S3).

### Iso-Seq based annotation of the IPO323 genome sequence and gene model selection

*Z. tritici* mRNAs used for this study corresponded to a wide array of *in vitro* mycelial growth conditions (Table S3). These mRNAs were used for the construction of either single-stranded Iso-Seq cDNA libraries or single-stranded Illumina cDNA libraries. The Iso-Seq sequences from each library were processed individually (cleaning, assembly) and pooled into a single dataset. Non-redundant Iso-Seq transcripts were selected at each locus using the CupCake chaining tool, giving 22,659 Iso-Seq transcripts. Some Iso-Seq transcripts corresponded to alternative transcripts differing in their intron splicing or TSS/TTS (TSS: transcriptional starting site, TTS: transcriptional termination site). The alternative Iso-Seq transcripts that were either not supported by RNA-Seq or with a relative abundance lower than 10% according to RNA-Seq in all conditions, were filtered out. This filtering kept isoforms differentially expressed in a least one condition with a relative abundance over 10%, providing 21,052 transcripts corresponding to 8,927 loci. Most loci displayed only one isoform (50%), while other loci had either 2 to 5 isoforms (42%), or at least 6 isoforms (8%).

Each single-stranded RNA-Seq library generated in the framework of this study and publicly available datasets (Table S3) were assembled separately and transcripts with weak expression levels (TPM<1) were removed. Between 8,600 and 13,000 filtered transcripts were obtained depending on the library and kept as a separate dataset providing 498,010 single-stranded assembled RNA-Seq transcripts as evidence. Most existing *ab initio* gene prediction tools use RNA-Seq assembled transcripts as evidence to infer the structure of gene models. However, currently only a few gene prediction tools (Eugene [10], LoReAn [47]) can use Iso-Seq transcripts as evidence. These two softwares were used to annotate the IPO323 genome sequence with Iso-Seq transcripts, RNA-Seq transcripts and reference fungal protein sequences as evidence. Eugene identified 15,810 gene models in the *Z. tritici* genome in a two-pass mode and strand-specific prediction allowing overlapping gene models on opposite strands. This number was reduced to 15,245 gene models after filtering out genes corresponding to TEs. LoReAn identified 11,537 gene models in the *Z. tritici* genome without overlapping gene models on the opposite strand, which were reduced to 11,497 after filtering out genes corresponding to TEs. Selection of the best gene model was performed with InGenAnnot using the novel Eugene and LoReAn gene predictions and the four existing ones (JGI, MPI, RRES, CURTIN). All these gene models were clustered into 17,147 metagenes.

For each comparison InGenAnnot computes an Annotation Edit Distance (AED) [48] that is a distance either between two gene models or between a gene model and an evidence. AED computing takes into account the number of overlapping bases, as previously described [48]. Two additional options were implemented in AED computation, such as a comparison limited to the CDS to avoid bias between annotations without or with UTRs (provided only by Eugene), and a penalty score of 0.25 on transcript AED scores in case of incongruence in splicing sites between transcript evidence and the gene model. Since it is difficult to compare AED values derived from protein evidence to those from transcript evidence, different AED scores were computed for each type of evidence. The gene models with the best AED scores with either transcript or protein evidence, or both types of evidence, were selected based on CDS comparisons. Gene models with an AED of 0.3 for transcript and/or an AED of 0.1 for protein evidence were selected (Figure 2). Gene models failing to pass the AED threshold, but predicted by at least four independent annotations, were retained to avoid the loss of gene models with low support from transcript or protein evidence (upper right square in Figure 2 corresponding to 1,846 gene models). These rescued genes models were mostly not conserved across fungi (upper right red bar in Figure 2) and frequently had low transcriptional support (upper green bar in Figure 2). For gene models overlapping on opposite strands, only the gene model with the best AED score was selected. Finally, 97 additional effector-encoding genes were predicted with the *rescue_effector* tool of InGenAnnot.

**Figure 2.**
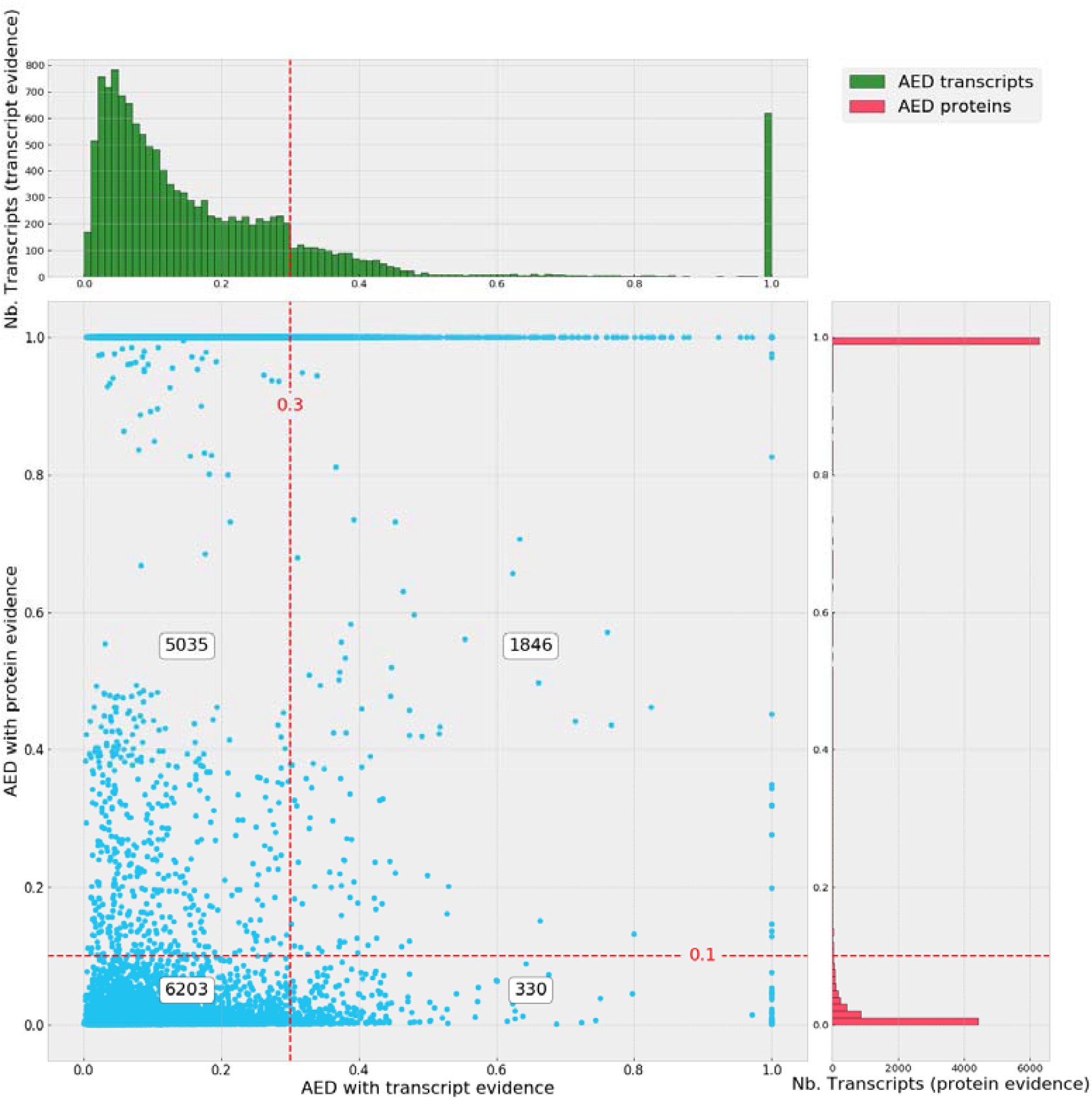
Selection of the best Re-annotated Gene Models (RGMs) according to their Annotation Edit Distance (AED) scores. Plot of RGM AED scores. AED scores (0-1) describe how a given gene model fits to transcript and protein evidence (best fit = 0). Transcript evidence was computed from RNA-Seq or Iso-Seq data (X axis). Protein evidence was computed from fungal protein sequences excluding *Zymoseptoria* species (Y axis). The red, dashed lines represent the AED thresholds to filter out genes (0.3 for transcripts, 0.1 for proteins), except if they are supported by at least four different annotations (1846 RGMs, upper right area of the graph). The numbers of genes in the four areas are displayed in white text boxes. Numbers of transcripts with transcript evidence were plotted on cumulative histograms above the scatter plot (green). Numbers of transcripts with protein evidence were plotted on cumulative histograms on the right of the scatter plot (red).

Overall, we obtained a final set of 13,414 re-annotated Gene Models (RGMs; File S1, Table S4). In addition, UTRs were inferred from Iso-Seq transcripts for 7,713 genes, and for 9,856 genes (73%) when combined with RNA-Seq assembled transcripts. The average and median sizes of 5’UTRs were 315 bp and 156 bp, while they were 389 bp and 220 bp for 3’UTRs (Table S4), close to the values (mean 5’UTR 275 bp and mean 3’ UTR 303 bp) reported recently for the Pezizomycotina *P. anserina* [70]. A small proportion of genes displayed long 5’UTRs (1 Kbp to 7 Kbp, 6%), and/or long 3’UTRs (1 Kbp to 8.6 kbp, 8.6%).

### Comparison of the reannotated IPO323 gene models with available genome annotations

The 13,414 IPO323 RGMs were compared to gene models predicted by the four previous annotations (JGI, MPI, RRES, CURTIN). This comparison was first performed using BUSCO and the ascomycota_odb as reference genes [49]. Higher BUSCO scores (99.4 % identical) were obtained with RGMs compared to the JGI, MPI and CURTIN annotations (95.7-98.5% identical), while scores obtained with RRES gene models were similar (99.1 % identical; Table S5). In particular, the JGI annotation had a high number of fragmented and missing BUSCO genes compared to other annotations, while the CURTIN annotation had a higher level of duplicated BUSCO genes compared to other annotations (Table S5). The eight missing BUSCOs in RGMs were reduced to six after manual inspection. These six RGMs that were missing in BUSCO encoded a Leucyl-tRNA synthetase, a WD40-repeat-containing domain protein, a Zinc finger protein, a Heavy metal-associated domain protein, a protein with an HMA domain, a PHD-type protein and a GTP binding domain protein. Their conservation across fungi is questionable, since a blastp search showed that they are missing from numerous genomes.

The comparison between annotations was then performed using AED scores (Figure 2, S1 and S2). Of the 13,414 RGMs, 11,568 (86%) passed the AED threshold of 0.3 and 0.1 for transcript and protein evidence, respectively (Figure 2). In comparison, these numbers decrease to 7,730, 8,936, 9,518 and 10,716 for the JGI, MPI, RRES and CURTIN annotations, respectively (Figure S1). This comparison showed that RGMs had a higher level of evidence support, followed by the CURTIN annotation, while JGI was the least-supported annotation. Among the 1,846 RGMs failing to pass the AED threshold, but rescued as predicted by at least four annotations, 574 have no AED score. This implied that they were only predicted by *ab-initio* software (see genes with no evidence in Table S6). 224 of these 574 fully *ab-initio* RGMs (40%) were located on the 3’ arm of chromosome 7 between positions 1,900,000 and 2,500,000 (Table S6). Almost none of these RGMs was expressed, even during infection. This region was previously described as carrying a high level of histone H3K27me3 and H3K9me3 modifications mediating transcriptional silencing, similar to those found in accessory chromosomes [71]. These marks could explain the lack of expression of genes from this region of chromosome 7. In addition, none of these genes was conserved across fungi, suggesting either a recent origin or an artefact from annotation pipelines. The other fully *ab-initio* RGMs were more frequently localized on accessory chromosomes (32-53%) than on core chromosomes (12-16%, Table S6).

Among the 13,414 RGMs, 7,888 were identical to at least one gene model from another annotation (Figure 3), while 3,479 RGMs were identical to all the gene models from the four previous annotations (Figure 3). Since 3,618 gene models were identical among the four previous annotations (see above), 139 of these genes were not identical to RGMs. Most of the corresponding 139 RGMs had a novel start codon that did not change the coding phase of the first open reading frame, leading to a shorter or longer version of the same protein compared to other annotations. However, these novel start codons were not necessarily more supported by transcript evidence than those from previous annotations. Ribosome profiling could help in solving this problem by identifying the real start codon [72]. 2,047 RGMs either differed from all gene models of other annotations (1,376, Table S6) or were not predicted by any other annotation (671, specific RGMs, Table S6). Most of the 1,376 RGMs differing from all other annotations had either alternative ATGs (see above) or intron splice sites supported by transcript evidence. RGMs also included novel gene models resulting from resolving the structure of incorrectly fused collinear gene models (see below).

**Figure 3.**
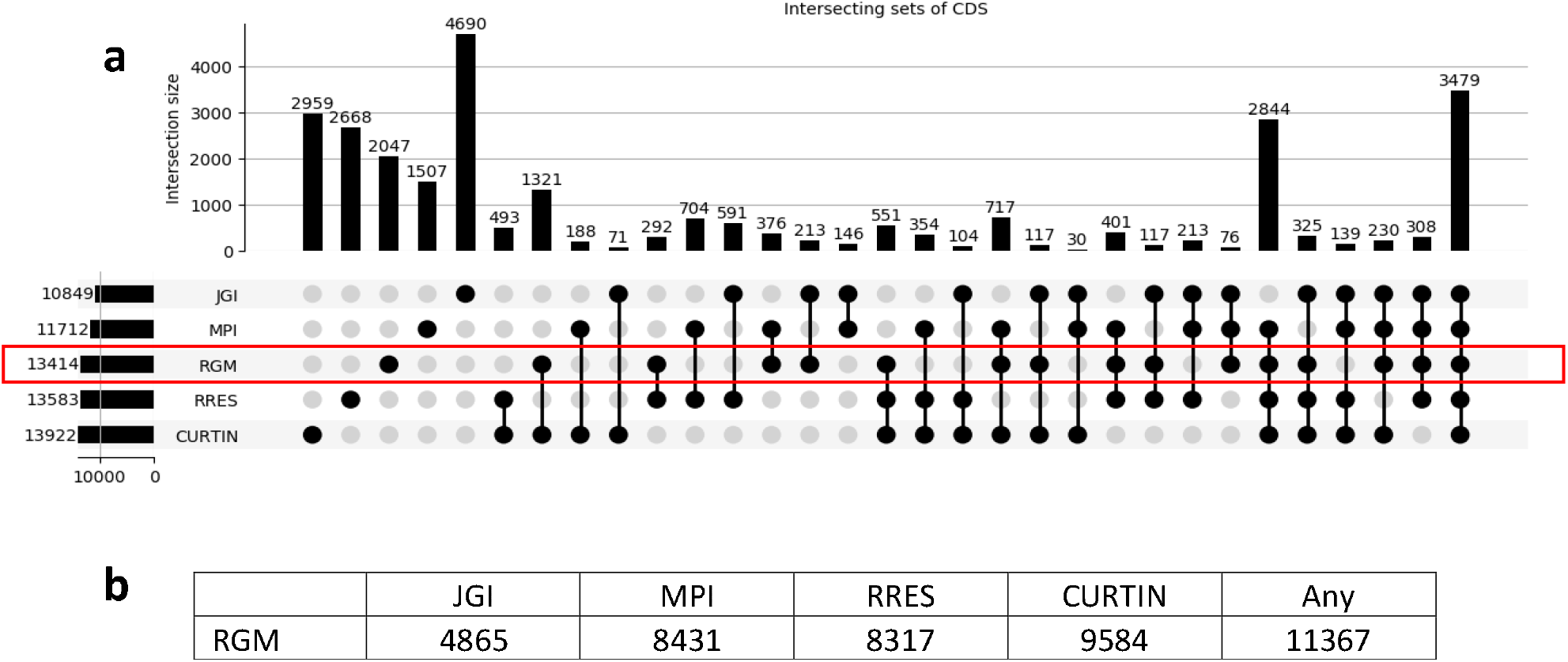
Comparison of the novel IPO323 genome annotation (Re-annotated Gene Models, RGM) with the four available annotations a) Upsetplot of RGMs with gene models from the four available annotations (JGI, MPI, RRES and CURTIN). Number of shared (identical) gene models for coding sequences (CDS). b) Number of identical CDS between RGMs and each available annotation.

The 671 specific RGMs were distributed evenly on all chromosomes (Table S6). 117 of these specific RGMs displayed more than 40% similarity to proteins from other fungi, including 63 with more than 80% similarity. A *tblastn* search against the 31 existing *Zymoseptoria* spp. genome sequences was performed. Most RGM specific genes were found in other *Z. tritici* strains (File S1), in particular in the genome of strain ST99CH_1A5 (571 hits with a at least 75 % identity and 75% coverage), while only a few hits were found in the most distant species *Z. passerinii* SP63 (22 hits). Overall, 654 of the 671 RGM specific genes (97%) matched at least one *Zymoseptoria* spp. sequence. These new genes were often located in regions with complex patterns of expression. A manual curation of these gene models will be required to confirm their accuracy.

One major improvement of RGMs was in resolving the structure of genes that were incorrectly fused in previous annotations (split RGMs). These genes were identified by detecting overlaps between gene models from different annotations. This survey revealed a high number of RGMs resulting from the splitting of fused genes from the MPI and JGI annotations (1,507 and 1,258, respectively, Table S7), and to a lesser extent from the RRES annotation (701), while these genes were in low number in the CURTIN annotation (176). The average AED score of split RGMs was better (median AED score: 0.17) than that of the fused gene models (median AED score: 0.34). In addition, most MPI fused genes (87%) were not supported by transcript evidence, since their AED scores were higher than the cutoff value (>0.3, Figure S3). On the reverse, most transcript AED scores of split RGMs (65 %) were supported by transcript evidence, since their AED scores were lower than the cutoff value (0.3<, Figure S3). Still, a significant number of split RGMs (494, 35%) had low support from both transcript and protein evidence (upper right square in Figure S3). These split RGMs were rescued since they were also identified in other annotations than MPI.

Overall, these results showed that the split RGMs were better supported by transcript and protein evidence than the MPI fused genes. The transcript evidence of two randomly chosen MPI fused genes and their corresponding split RGMs is shown in Figures S4 and S5. Both MPI fused genes had no Iso-Seq transcript support, while Iso-Seq transcripts supported the corresponding split RGMs. Assembled RNA-Seq transcripts supporting split RGMs were also observed for RGM-1 and RGM-2 from Figure S4. However, large assembled RNA-Seq transcripts were supporting the fused MPI gene model from Figure 5. Still, some of these assembled transcripts included alternative introns that were not supported quantitatively by RNA-seq. We hypothesise that these long, chimeric transcripts were artefacts of the assembly of RNA-Seq reads from individual genes with overlapping transcripts. The final proof supporting these split RGMs was obtained by identifying specific expression conditions (13 days post-inoculation, wheat infection, Figure S5) in which RGM-2 was strongly expressed, but not RGM-1.

### Functional annotation of the reannotated IPO323 gene models

Functional annotation of predicted proteins deduced from RGMs was performed using both Blast2Go and InterProScan. 5,593 RGMs exhibited a GO term or an IPR and 2,838 were annotated with at least one Enzyme Code (EC). As in previous annotations of IPO323 genome sequence [28], [73], several tools were launched to identify genes encoding putative secreted proteins, including effectors (File S1). We identified 1,895 genes corresponding to secreted proteins with less stringent criteria than those used in a previous study that identified 970 secreted proteins using the JGI annotation [43]. All these 970 genes were identified as RGMs. However, they increased to 1,046 mainly due to the splitting of fused gene models from the JGI annotation. The RGM secretome included 234 small, secreted proteins (SSP) according to EffectorP and additional criteria defined in the Materials and Methods section. Among the 100 SSPs studied previously by Gohari et al. using the JGI annotation [74], 93 were identified as encoded by RGMs. Still, many structural differences between these RGMs and the JGI gene models were observed. The effector rescue software of InGeAnnot identified 53 SSPs among which 43 were not found in any previous annotations. Four of these 53 novel SSPs displayed a significant upregulation during infection compared to *in vitro* culture conditions (ZtIPO323_001210, ZtIPO323_072700, ZtIPO323_105940 and ZtIPO323_123970), suggesting a possible role in infection. In addition, genes encoding effectors missing in previous annotations, such as *Avr-Stb6*, were now predicted correctly. The new annotation also predicted two additional *Avr-Stb6* paralogs located on chromosome 10 (Figure S6a), while the original *Avr-Stb6* is located at the end of chromosome 5 (Figure S6b, [32]).

### Identification of alternative transcripts using combined Iso-Seq and RNA-Seq evidence

The initial set of 21,052 Iso-Seq transcripts used for gene reannotation was filtered to exclude UTR length isoforms, yielding 11,690 Iso-Seq transcripts corresponding to coding and non-coding loci. Sqanti3 allocated 10,938 Iso-Seq transcripts to 8,199 RGMs (Table 1). 7,872 of these RGMs had the same structure as their matching Iso-seq transcripts (full_splice_match). The other 327 RGMs, classified as *“*ISM*”* or *“*genic*”* by Sqanti3 displayed a structure differing from their matching Iso-seq transcripts. These gene models were supported either by other evidences (RNA-Seq, protein) or rescued (*ab initio* only). In most cases, these Iso-Seq transcripts were only partly covering the RGMs, suggesting that they were partial cDNAs likely due to the early termination of reverse transcription. 2,716 Iso-Seq transcripts were identified as alternative splice variants (25 % of coding transcripts). They were classified by Squanti3 into the following events: combination of known splicing sites (NIC); new splicing sites (NNC); intron retention (IR); and genic (Table 1). Most alternative transcripts corresponded to intron retention events (IR, 75%). Since transcripts could carry a premature termination codon (PTC) recognized by the non-sense mediated decay (NMD) pathway, they were screened for potential NMD signals [75], leaving 2,372 alternative transcripts corresponding to 1,742 RGMs. The numbers of RGMs with 2, 3, 4 and at least 5 isoforms were 1,342, 274, 77 and 49, respectively (Table S8). A total of 337 alternative transcripts corresponded to a novel assembly of coding exons, 271 to a novel assembly of UTR exons, and 16 to a novel assembly of both (included in NIC, NNC and Genic events, Table 1). For example, RGM ZtIPO323_030030, predicted to encode a putative SSP in a previous study (SSP10, [76]), had an alternative splicing site providing a new exon and a shorter protein that was reduced by 34% in length at its C-terminus (Figure 4a). The 1,753 remaining isoforms with intron-retention events could correspond to un-spliced transcripts not detected by our NMD screen. Some alternative transcripts were detected in high amounts by RNA-Seq, as observed for RGM ZtIPO323_013330 (Figure 4b) with two intron-retention events. This RGM has 4 transcript isoforms. The canonical transcript (Iso-Seq 2), corresponding to the structure of the selected RGM, had 4 splicing sites, one being located in the 5’ UTR. Two alternative Iso-Seq transcripts (Iso-Seq 1 and 2) with one or two intron-retention events were also supported by RNA-Seq. The last Iso-Seq transcript (n°4) had an alternative splicing of the fourth intron that was not supported by RNA-Seq data. Some alternative transcript isoforms were used as a major evidence for selecting the RGM as shown for ZtIPO323_030030 (Figure 4a) or ZtIPO323_013090 (Figure S7). These examples illustrated the difficulty for gene predictors to choose between gene models with complex alternative splicing events or co-existing isoforms with similar expression levels (Figure 4a).

**Table 1.** Classification of Iso-Seq transcript isoforms from Zymoseptoria tritici isolate IPO323 Filtered Iso-Seq transcripts from different growth conditions were analysed and classified with Sqanti3.

| Categories | Counts |
| --- | --- |
| Full-splice_match (FSM) <sup>1</sup> | 7872 |
| Incomplete-splice_match (ISM)<sup>2</sup> | 305 |
| Fusion | 45 |
| Genic<sup>3</sup> | 664 |
| Intron retention (IR) | 1571 |
| novel_in_catalog (NIC)<sup>4</sup> | 7 |
| novel_not_in_catalog (NNC)<sup>5</sup> | 474 |
| Antisense | 395 |
| Intergenic | 357 |

<sup>1</sup> Whole transcripts with possible alternative 3' and 5' ends
<sup>2</sup> Partial overlaps of transcripts fitting with intron coordinates
<sup>3</sup> Partial overlaps of introns and exons not compliant with intron/exon coordinates
<sup>4</sup> Use combination\_of\_known\_splice sites
<sup>5</sup> At\_least\_one\_novel\_splice site detected

**Figure 4.**
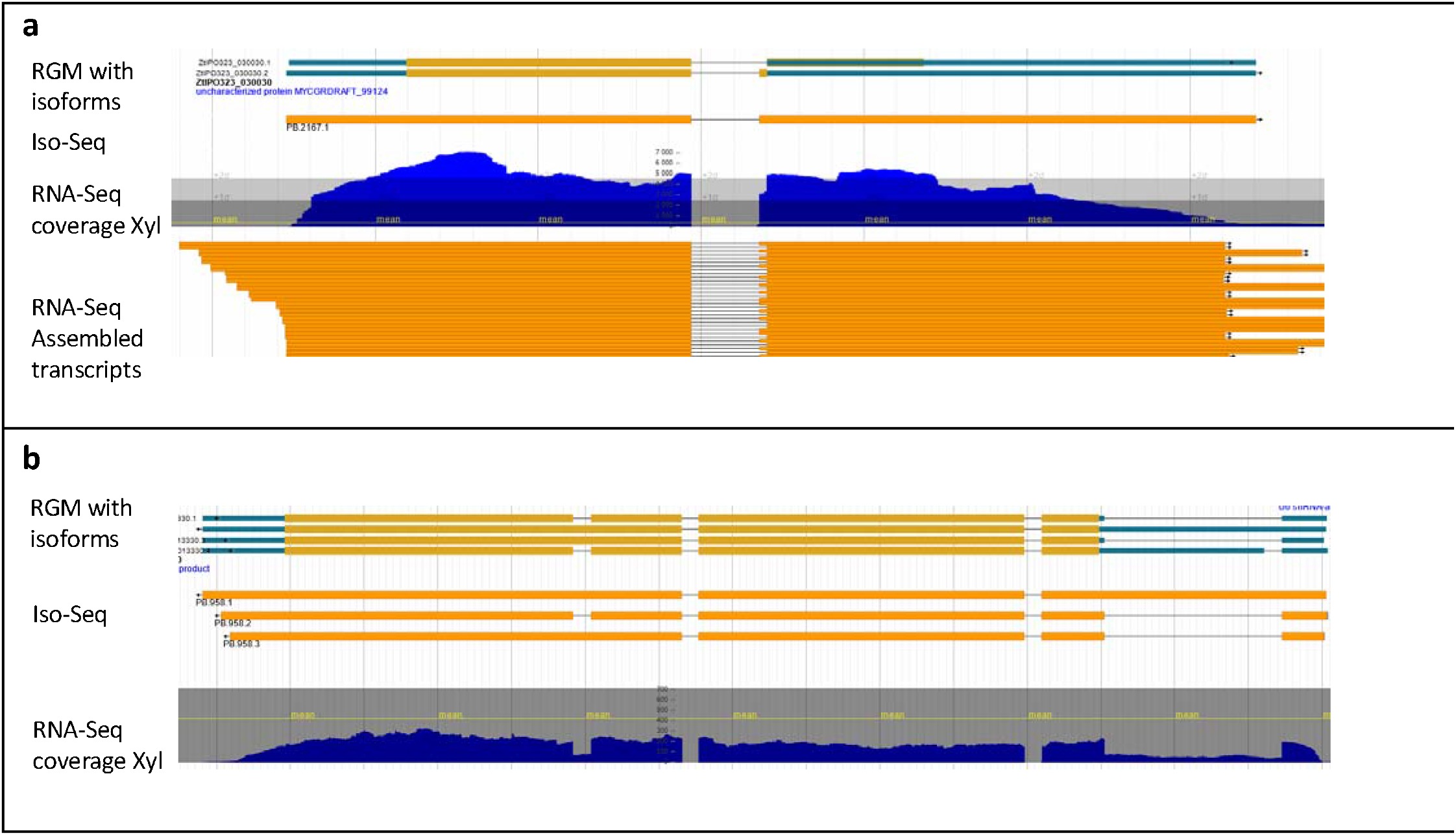
Transcript isoforms of Re-annotated Gene Models (RGMs) ZtIPO323_030030 (a) and ZtIPO323_013330 (b) supported by Iso-Seq and RNA-Seq evidence. a) Gene ZtIPO323_030030 (chr2: 777930 … 1778675, 747 b). This RGM has two transcript isoforms (alternative 3’ acceptor site). Both encoded Small Secreted Proteins (SSP 10, File S1). Previous annotations selected the second acceptor site leading to the longest CDS. A single Iso-Seq transcript corresponding to the longest CDS was detected (Iso-Seq track), while both isoforms were detected using RNA-Seq data (RNA-Seq assembled transcript). RNA-seq coverage identified both isoforms in equal amounts (RNA-Seq coverage Xyl). Based on read coverage from different RNA-Seq libraries, the isoform corresponding to the shortest CDS was the most frequent. This isoform was likely the canonical form and encoded a protein with a C-terminus that was reduced in length by 34% compared to the other isoform. RGMs with isoforms track: different isoforms. Iso-Seq track: filtered Iso-Seq transcripts. RNA-Seq coverage Xyl track: coverage of strand-specific RNA-Seq reads. RNA-Seq assembled transcript track: assembly of strand-specific RNA-Seq reads. b) ZtIPO323_013330 (chr_1:3420115..3424093, 3.98 Kb). This RGM had four transcript isoforms. The selected RGM had four splicing sites, one of which in the 5’ UTR was supported by Iso-Seq transcript (Iso-Seq n°2) and RNA-Seq (RNA-Seq coverage Xyl). Two Iso-Seq transcripts with one or two intron retention events were detected as Iso-Seq transcripts (Iso-Seq n°1 and 3) and confirmed by RNA-Seq (RNA-Seq coverage Xyl). One Iso-Seq transcript had an alternative 5’ donor splicing site in the 5’ UTR (Iso-Seq n°4). This isoform was likely weakly expressed, as it was not supported by RNA-Seq (RNA-Seq coverage Xyl). RGMs with isoforms track: different RGM isoforms. Iso-seq track: filtered Iso-seq transcripts. RNA-Seq coverage Xyl track: coverage of strand-specific RNA-Seq reads. RNA-Seq assembled transcript track: assembly of strand-specific RNA-Seq reads.

### Differential expression of Iso-Seq transcript isoforms

RNA-Seq data were used to detect differential isoform usage (DIU) for coding genes. RGMs with significant DIU between different *in vitro* culture conditions or between infection and *in vitro* culture conditions were identified using tappAS [29] with a minimal p-value of 0.01. Only 22 RGMs had a DIU between different culture conditions, in particular between Galactose/Sucrose and Mannose/Xylose growth media (File S1). Ten of them were associated with GO terms (GTPase activity, ATP and GTP binding). A total of 163 RGMs displayed a DIU between at least one infection time point and one culture condition, and 88 (54%) encoded proteins with GO terms (File S1), including 23 secreted proteins. The number of these genes was too small to perform a GO enrichment test. 30 of these 163 RGMs were specifically up or down regulated during infection compared to all culture conditions including ZtIPO323_042160 and ZtIPO323_042360, encoding proteins without known function, and ZtIPO323_043800, encoding a PHD and RING finger domains-containing protein. Two of these 30 DIU genes (ZtIPO323_016670 and ZtIPO323_043500) encoded secreted proteins that were significantly upregulated at late infection stages (13, 21 dpi). ZtIPO323_016670 encoded a carbohydrate esterase from family 8 involved in cell wall modifications and ZtIPO323_043500 encoded a SSP. Manual inspection of the RNA-Seq data associated with these DIU RGMs confirmed their differential expression, but not a different usage of isoforms. Indeed, the isoforms detected during infection corresponded to a low number of reads compared to *in vitro* culture conditions. This could lead to a bias in DIU analyses.

### Identification of long non-coding RNAs and survey of their expression

Sqanti3 allocated 752 Iso-Seq transcripts to non-coding loci (Table 1). Among these transcripts, we identified 395 antisense and 357 intergenic non-coding transcripts. These 752 Iso-seq transcripts were analyzed for the presence of long non-coding RNAs (lncRNAs). Most previous analyses of fungal lncRNAs were performed using RNA-Seq data with a 200 bp minimal size cutoff. A single study of fungal lncRNAs was performed using Iso-seq in *F. graminearum* [77]. This study showed that lncRNAs were generally larger in size than 1 kb. Therefore, we chose a cutoff value of 1 kb in length for selecting candidate lncRNAs. *Z. tritici* Iso-seq transcripts overlapping with TEs, smaller than 1 kb in length and containing an ORF longer than 300 bp (100 amino acids) were discarded. Changing the 1-kb length threshold to 200 bp only removed 72 lncRNAs. This selection left 398 candidate lncRNAs (288 antisense and 110 intergenic). As previously observed [77], intergenic lncRNAs are generally smaller than antisense lncRNAs, explaining the strong impact of size selection on this category. Filtering ORFs longer than 300 bp removed 343 lncRNAs, representing a large proportion of the 398 candidate lncRNAs (86%). We decided to keep this stringent criterion to select only reliable lncRNAs. This criterion avoided selecting lncRNAs encoding coding genes not retained by InGenAnnot. For example, the Iso-Seq PB.5809.X located on chromosome 7 (position 688635 to 690776 bp), for which Eugene predicted a gene model not retained as an RGM, was removed from candidate lncRNAs using this criterion. This process selected 55 lncRNAs, among which 3 were labelled as “coding” based on their coding potential and 1 contained an ORF with a pfam domain. Finally, 51 transcripts were classified as lncRNAs according to our stringent criteria and 35 of these lncRNAs (68%) were differentially expressed in at least one pairwise comparison (p-value 0.05). Half of these lncRNAs were differentially expressed between infection and *in vitro* growth conditions, including 5 that were up-regulated and 12 down-regulated during infection (log2FC > 2). Most lncRNAs that were down-regulated during infection were antisense transcripts (83%). The lncRNA PB1188.1 was down-regulated during infection compared to all culture conditions (Table S9). This lncRNA was an antisense transcript of ZtIPO323_016330, encoding a secreted Subtilisin-like protein, that was up regulated during infection but down regulated during *in vitro* culture conditions. Another RGM (ZtIPO323_037670) encoding a TTL protein (Tubulin tyrosine ligase involved in the posttranslational modification of tubulin) and its antisense lncRNA PB.2709.1 displayed a negative correlation with their expression pattern during infection (Table S9). In this case, the antisense lncRNA PB.2709.1 was up regulated during infection, while the corresponding coding gene ZtIPO323_037670 was down regulated.

### Iso-Seq transcripts revealed polycistronic mRNAs

Alignment of Iso-Seq transcripts with RGMs identified 2,625 potential polycistronic transcripts. Among them, 224 corresponded to polycistronic transcripts containing two to three RGMs on the same strand supported by independent long-read single-transcript molecules. For example, adjacent RGMs ZtIPO323_010430 and ZtIPO323_010440 were transcribed on the same strand with overlapping 3’UTR and 5’UTR (Figure 5, red rectangle). Iso-Seq polycistronic single-transcript molecules covering the two RGMs were detected, as well as single RGM Iso-Seq transcripts (Figure 5, Iso-Seq track and Iso-Seq polycistronic track). Assembled RNA-Seq reads at this locus mostly predicted a transcript covering the two RGMs (Figure 5, RNA-Seq transcripts tracks). This long transcript likely resulted from the wrong assembly of reads from overlapping transcripts. Indeed, RNA-Seq coverage strongly decreased in the region of the overlap between the two RGMs, suggesting two independent transcripts (Figure 5, RNA-seq coverage track). This RNA-seq coverage analysis also suggested that the abundance of the polycistronic transcript was low compared to single-gene transcripts. Multiple stop codons were present in these polycistronic transcripts, excluding the possibility of errors in annotated genes for a larger single ORF, as observed for polycistronic transcripts described in *Agaricomycetes* [78], and *F. graminearum* [77] or *Cordyceps militaris* [79].

**Figure 5.**
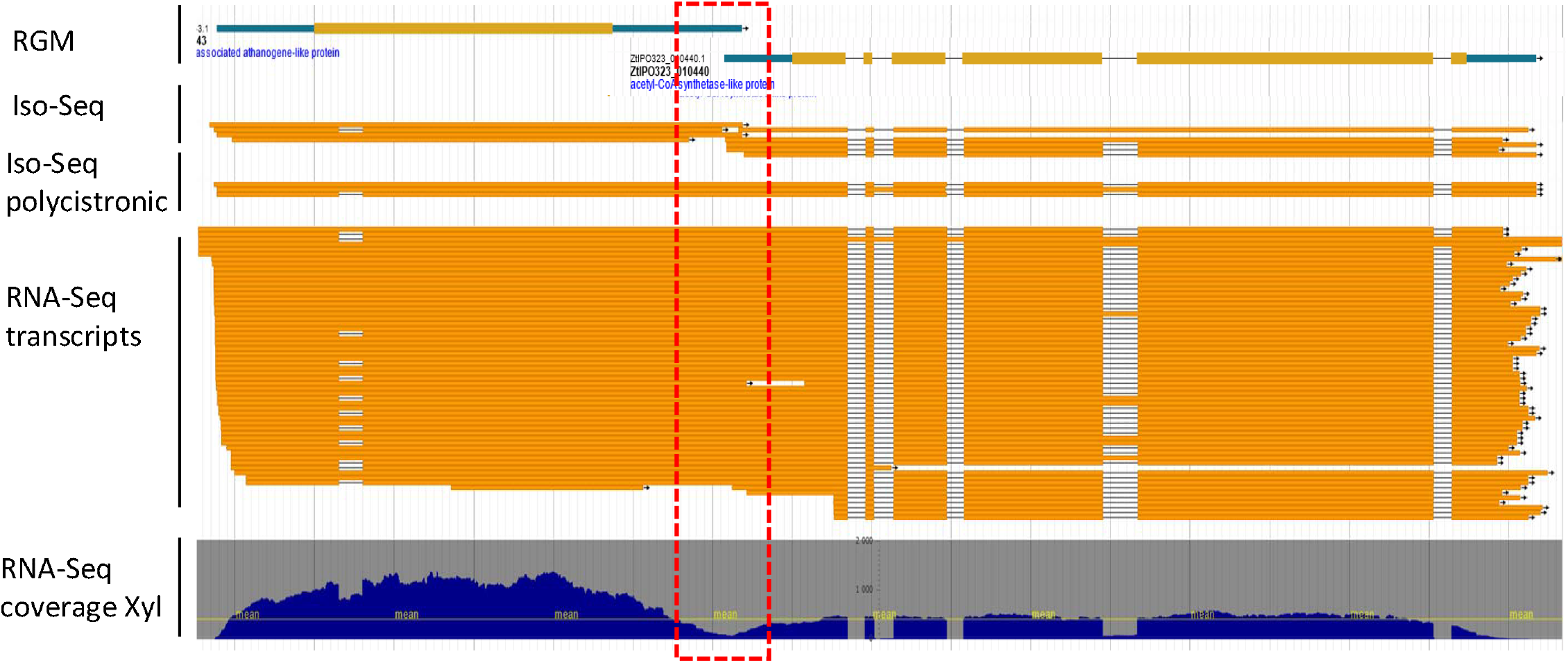
Examples of polycistronic transcripts shown for Re-annotated Gene Models (RGMs) ZtIPO323_010430 and ZtIPO323_010440 RGMs ZtIPO323_010430 and ZtIPO323_010440, located at chr_1:2692858…2697168 and chr_1:2692858…2697168, respectively, were transcribed on the same strand with overlapping 3’UTR and 5’UTR (red rectangle). Iso-Seq polycistronic track: evidence of transcripts covering the two RGMs. A strong decrease in RNA-Seq coverage was observed in the region of the overlap (red dashed rectangle), suggesting two singles, overlapping transcripts. The assembly of RNA-Seq reads led to a polycistronic transcript involving the two RGMs, likely resulting from the wrong assembly of reads from these overlapping transcripts. Iso-seq track: filtered Iso-seq transcripts mapping at this locus. Iso-Seq polycistronic track: polycistronic transcripts identified in the Iso-Seq database. RNA-seq transcript track: assembly of strand-specific RNA-Seq reads mapping at this locus. RNA-seq coverage Xyl track: coverage of strand-specific RNA-Seq reads mapping at this locus.

### Iso-Seq transcripts encoding fungal mycoviruses

A total of 2,203 Iso-Seq transcripts did not map to the *Z. tritici* IPO323 genome and were discarded for annotation. These transcripts were clustered and analysed for their similarity with known sequences. The larger cluster of independent Iso-Seq transcripts (1919 sequences) was identical to Fusarivirus 1 (ZtFV1), already identified by a large-scale fungal transcript analysis [80]. The second cluster gathered 17 independent Iso-Seq transcripts that were closely related to narnavirus 4 of *Sclerotinia sclerotiorum* (SsNV4) [81]. As these viral Iso-Seq transcripts were probably obtained by internal polyA priming, they did not cover the full sequence of the viruses. To rescue the full-length viral RNA, *de novo*-assembly was performed using RNA-Seq data mapping to the viral Iso-Seq consensus sequences. RNA-Seq reads corresponding to these two fungal viruses were detected in all our cDNA libraries. These analyses showed that the ZtFV1 Iso-seq transcript was a full-length viral sequence. However, the second viral Iso-Seq transcript related to SsNV4 was shorter than the viral RNA assembled from RNA-Seq reads. This allowed the reconstruction of a full sequence of 3091 nucleotides encoding a protein of 986 amino acids corresponding to a RNA-dependent RNA polymerase. This new virus, ZtNV1 (*Zymoseptoria tritici* NarnaVirus 1), is as long as SsNV4 (3105bp). ZtNV1 displayed 71% identity at the nucleotide level and 67% identity (79% similarity) at the protein level with SsNV4. The phylogenetic tree of viral RNA-dependent RNA polymerases showed that the ZtNV1 was highly related to narnaviruses from *S. sclerotiorum, Plasmopara viticola*, and Fusarium asiaticum (Figure S8). IPO323 ZtNV1 sequence was used to screen publicly available *Z. tritici* RNA-seq datasets. ZtNV1 was identified in all these datasets, but only with very few reads, validating the ubiquitous presence of the virus in *Z. tritici*. ZtFV1 was also detected in these RNA-seq data in higher amounts compared to ZtNV1 (70,000 fold).

## Discussion

### Improvement of the *Z. tritici* IPO323 gene models

We developed a new strategy to generate high-quality genome annotations using the fungus *Z. tritici* as a case study. The major requirement for improving the *Z. tritici* IPO323 genome annotation was the production of a set of full-length transcript sequences. Gene annotation strongly relies on transcriptomic data to support the structure of a predicted gene and define its boundaries. The assembly of RNA-Seq short reads frequently leads to artefacts such as chimeras corresponding to adjacent genes with overlapping transcripts [16], especially in genomes with a high gene density [37]. Iso-Seq long-read by-pass these artefacts, as it produces sequences from single cDNA molecules without assembly. Iso-Seq also provides transcript isoforms corresponding to alternative start, stop and splicing events. Still, Iso-seq has potential pitfalls since this technic is not quantitative. Indeed, we identified rare Iso-seq transcripts likely corresponding to errors of the transcriptional machinery (intron retention, polycistronic transcripts). We minimized this error by filtering out low-abundance Iso-Seq transcripts based on their quantification using short-read RNA-seq. Overall, filtered Iso-seq transcripts were highly reliable in determining the genome-aligned exon structure of transcripts, while RNA-Seq offered a quantification of Iso-Seq transcript structures and isoforms.

The newly established transcriptomic dataset was used to select the best gene models among those predicted by different *ab initio* software according to their AED transcript scores (transcript evidence), using InGenAnnot. Protein evidence also helped select the best gene model for genes not expressed under the conditions used for producing mRNAs. The combination of six *ab initio* software was needed at two levels. First, a diversity of software was needed to produce a sufficient number of gene models at each locus to be selected by InGenAnnot. Indeed, none of the *ab initio* software was able to independently predict all the RGMs (Table S10). The best *ab initio* software, Eugene, only predicted correctly 76% of the RGMs. Second, the use of different *ab initio* software allowed the rescue of gene models without evidence (1,846 RGMs predicted by at least 4 different *ab initio* software). Most rescued RGMs were not conserved across fungi and they had a low transcriptional support or they were not expressed under the available conditions (upper green bar in Figure 2). They typically included candidate fungal effectors that could be important for plant-fungal interactions (File 1). Yet, these rescued RGMs may be artefacts of *ab initio* software, and they need to be validated manually.

Overall, our strategy significantly improved the annotation of the *Z. tritici* IPO323 genome, and missing genes encoding effectors such as Avr-Stb6 were now predicted correctly. In addition, it revealed different bias in previous annotations. Among the 13,414 RGMs, 2,047 were either different from all previous gene models (1,376, Table S6) or not predicted in previous annotations (671 RGM-specific, Table S6). We are confident that changing/adding these RGMs is an improvement in the prediction as both transcripts and protein evidence supported these changes. The most frequent discrepancy was the occurrence of fused genes in previous annotations that were split into distinct RGMs. Most of these fused genes corresponded to RGMs with overlapping transcripts (Figures S4, S5). Indeed, the assembly of RNA-Seq reads corresponding to such transcripts could have generated chimeric transcripts, providing erroneous evidence to the software used in these annotations. Changes in parameters used for RNA-Seq read assembly could reduce the number of chimeric transcripts. However, Iso-Seq long-read sequencing clearly avoided this artefact and its use as transcript evidence likely explains the observed improvement in the RGMs. To our knowledge, only two previous studies improved fungal gene prediction using Iso-Seq transcript long-read sequences (*C. militaris*, [79]; *F. graminearum*, [77]). We further improved the method used in these papers by filtering Iso-Seq transcripts according to their abundance, and by creating a method to select the best gene model according to different *ab initio* annotations and evidence.

### Iso-Seq long reads reveals the complexity of transcripts in Z.tritici

Identifying transcript isoforms is a major challenge when relying on the assembly of short RNA-seq reads, as alternative splicing sites could not be easily distinguished. Here, we took advantage of the full-length cDNAs produced by Iso-Seq long-read sequencing to identify novel exon combinations. Indeed, the assembly of RNA-Seq reads could be misleading for transcripts with more than one. However, Iso-Seq sequencing is not a quantitative method and minor transcripts were sequenced. For example, Iso-Seq transcript isoforms with long UTRs or IR without strong support from RNA-Seq data were identified in our initial dataset (Figure 4, Figure 5, Figure S5). These low-abundance transcript isoforms could be produced by the transcriptional machinery either as by-products or to regulate gene expression. As observed for gene annotation (see before), the best strategy is to filter Iso-Seq sequences with RNA-Seq data to withdraw transcript isoforms with weak quantitative support, with the caveat that some transcripts might be excluded. As observed in other fungal genomes ([77], [82], and references quoted within), most alternative splicing events were intron retention (IR). Indeed, we identified 58% of alternative transcripts with IR after NMD filtering (Table 1). IR events could generate premature termination codons (PTCs) likely degraded by the NMD pathway. However, NMD signals are difficult to predict with current bioinformatics tools in filamentous fungi. DIU analysis revealed a few RGMs with differential expressed transcript isoforms during infection compared to *in vitro* growth conditions. As discussed before, the small amounts of RNA-Seq reads available in these conditions makes such comparisons difficult using the available statistical tools. In fact, manual inspection of several detected loci did not reveal clear patterns of DIU for alternative transcripts.

Additionally, dense genomes, such as *Z*.*tritici* genome, are suitable for polycistronic transcription, i.e. the production of mRNA that encode several proteins. Indeed, we identified polycistronic mRNAs in *Z. tritic* among Iso-Seq long-read transcripts, as already observed in *Agaromycotina* [78] and *F. graminearum* [77] or *C. militaris* [79] using Iso-Seq. However, polycistronic-specific RNA-Seq reads were always detected in low abundance compared to single-gene transcripts. These RNA-seq data also showed that polycistronic transcripts mostly corresponded to genes with transcripts overlapping those from adjacent genes. As Iso-Seq is sensitive enough to detect rare transcripts, it is possible that these polycistronic transcripts are rare read-through transcripts. This hypothesis is supported by the fact that *in vitro* culture conditions of yeast known to be associated with increased transcriptional read-through led to more polycistronic transcripts [83]. Alternatively, these polycistronic transcripts could be an additional level of transcriptional control.

### lncRNAs are differentially expressed during wheat infection

LncRNAs are important components of transcriptional and translational regulation [84]. They can act in *cis* or *trans* of target genes, and modulate their expression by different mechanisms, leading to either the up-regulation or down-regulation of target genes [84]. Most of studies on fungal lncRNAs used assembled RNA-Seq reads [85]. This approach could lead to assembly artefacts. Iso-Seq long reads bypass this problem as entire cDNA molecules were independently sequenced. This process facilitated the identification of full length, non-chimeric lncRNAs. Using stringent criteria (size > 1000 bp, no ORF > 100 aa, no overlap with TEs), we identified 51 lncRNAs in *Z. tritici*. This number is far lower than those identified in other fungi (939 in *N. crassa* [86], 352 in *Verticillium dahliae* [87], and 427-819 in *F. graminearum* [77]). This difference could be due to the stringent criteria used for this study. In fact, when using similar criteria to previous studies, such as keeping all ORFs with no coding potential independently of their size, we identified 398 lncRNAs. In addition, many lncRNAs identified in these fungi were detected in specific conditions corresponding to stress [86], [88], and sexual development which we did not sample [77].

We investigated the role of lncRNAs in the wheat leaf infection by *Z*.*tritici*, and identified that 17 of the 51 lncRNAs were differentially expressed during plant infection, mostly as antisense transcripts (Table S9). Among them, two displayed expression patterns opposed to their corresponding coding genes. The lncRNA PB1188.1 was down-regulated during infection compared to *in vitro* culture conditions. This lncRNA is an antisense transcript of ZtIPO323_016330 encoding a secreted Subtilisin-like protein, that is up-regulated during infection. Subtilisin-like proteins are known to be secreted proteases playing an important role in plant infection [89], [90] and in plant–pathogen interactions [91], [92]. This negative correlation suggested that the down regulation of lncRNA PB1188.1 during infection allowed the full expression of ZtIPO323_016330 in infected leaves. The second lncRNA (lncRNA PB.2709.1) was up-regulated during infection compared to *in vitro* culture conditions (Table S8), while its corresponding transcript (ZtIPO323_037670) was down-regulated during infection. This transcript encodes a tubulin tyrosin ligase (TTL), a protein involved in the post-translational modification of tubulin. Thus, reduced expression of a TTL protein could alter tubulin turnover during infectious growth. The negative correlation observed between the gene expression and the expression of the corresponding antisense lncRNA suggests that antisense lncRNAs could be involved in the control of fungal gene expression during infection. Our observation hints at the existence of co-regulation networks between coding and non-coding transcripts in *Z. tritici* and suggest that this mode of regulation could be important during infection, as already observed during the infection of rice leaves by *M. oryzae*, [93]. These examples stress the importance of including lncRNAs in future studies to gather a comprehensive picture of the expression regulation landscape in *Z*.*tritici*.

### RNA mycoviruses are widespread in *Z*.*tritici*

In addition to the genes belonging to the *Z*.*tritici* genome, we revealed the presence of two RNA mycoviruses in IPO323. The first one Fusarivirus 1 (Zt-FV1) had been previously identified in *Z. tritici* by the screening of unmapped fungal RNA-seq reads [80]. We also identified a novel mycovirus, Zt-NV1 (Figure S8), related to the narnavirus 4 of *Sclerotinia sclerotiorum* (SsNV4) [81]. Using the Isoseq Zt-FV1 and Zt-NV1 sequences as templates, we retrieved RNA-seq reads corresponding to these mycoviruses in all of the IPO323 RNA-seq conditions tested, as well as from publicly available *Z. tritici* RNA-seq data, showing that these mycoviruses are widespread in *Z. tritici*. Zt-FV1 was the most abundant mycovirus, while Zt-NV1 was only detected as very few reads compared to Zt-FV1 (1/70,000), suggesting that it is a minor virus. Mycovirus are known to induce strong phenotypic defects in other fungi, so additional studies are needed to evaluate the role of these widespread mycoviruses in the life cycle of *Z. tritici*, in particular its growth, sporulation and pathogenicity [94].

### InGenAnot a novel tool for improving gene structure prediction

Many tools [8], [10]–[13], [95] and protocols [96] were established to predict gene models in eukaryotic genomes. Some were dedicated to fungal genome annotation [15], [35], [37] and were incorporated in bioinformatics workflows [14]. Evaluation of the reliability of an annotation is not an easy task. One of the most frequently used tools is the BUSCO software for identification of conserved proteins to evaluate the completeness and fragmentation of the predicted genes at the protein level [49]. More recently, new datasets and methods were proposed to test the reliability of gene annotations, looking deeper into the prediction of intron and exon structures [7]. However, this evaluation was still based on selected datasets, representing a conserved and partial view of gene content of a genome. In the case of a genome reannotation, ParsEval could give clues on overlaps of different versions of annotations with sensitivity and specificity metrics [69]. The most descriptive tool to evaluate the reliability of an annotation with associated evidence is GAEVAL (available through AEGeAn [97]), which computes an integrity score weighted by such features as confirmed introns, annotation coverage and UTR identifications.

In our new software, we implemented the AED metrics [48], to evaluate the ability of a gene structure to match with transcript evidence or other gene sets. We improved on previous implementation of the AED[11] by computing the AED metrics for each type of evidence (transcript and protein) and using a distinct score for Iso-Seq transcripts when available. Moreover, we allow penalized scores in case of discrepancy between the predicted structure and evidence, for example, when predicted splice sites were not supported. This evidence-driven annotation strategy required an in-depth analysis of data provided as evidence to eliminate potential artefacts. As each tool implements specific ML models, with different specificity/sensibility for each data source, their implementation and training parameters are more or less tolerant to particularities such as short CDS length or non-canonical splicing site. The combination of different gene prediction software with distinct intrinsic characteristics, could be a good way to avoid drawbacks from each software, in particular when *ab-initio* gene predictors fail to find a consensus gene model. In the same way as EvidenceModeler [36] or TSERBA [98], InGenAnnot is able to select the best gene model based on AED scores with defined evidence thresholds. We used additional criteria to select the best gene model when evidence was lacking (gene model predicted by all or a minimal number of software). Since each gene model had AED metrics, it could be compared to other gene sets, allowing post-filtering or prioritization in the manual curation process.

## Conclusion

In the era of the massive sequencing of compact fungal genomes, inferring gene models by evidence is essential and complementary to *ab-initio* gene prediction methods. In this paper, we used the recent Iso-seq technology and developed a novel software, InGenAnnot, to drastically improve the gene annotation of *Z*.*tritici*, an important fungal plant pathogen. We additionally identify lncRNA and mycoviruses as being expressed during plant infection. We expect that both the improved sequencing technology and our new software will be used widely to improve the gene prediction of many species of importance, in particular in plant pathogens with dense genomes, and reveal new insights into the role of transcriptome complexity in plant-pathogen interactions.

## Availability of data and materials availability

All raw sequencing data generated in this study have been submitted to the NCBI Gene Expression Omnibus (GEO) under accession GSE218898 with data accessions: GSM6758342 to GSM6758379. Processed data files of assembled RNA-Seq transcripts and filtered Iso-Seq reads were associated to the submission. Sequence of the new mycovirus ZtNV1 was deposited to NCBI under accession OP903463. Previous *Z. tritici* IP0323 gene annotations, new annotations (RGMs, Isoforms, LncRNAs) and annotation file, denoted file S1 (z.tritici.IP0323.annotations.txt), are available at: https://doi.org/10.57745/CVIRIB.

A genome browser with all annotations and evidence was set up at: https://bioinfo.bioger.inrae.fr/portal/genome-portal/12/ A new IPO323 genome web site at (https://mycocosm.jgi.doe.gov/Zymtr1/Zymtr1.home.html) was released with new genome annotations.

The InGenAnnot code and project is available at: https://forgemia.inra.fr/bioger/ingenannot Licensed under GNU GPL v3. InGenAnnot documentation is available at https://bioger.pages.mia.inra.fr/ingenannot

## Acknowledgments

We are grateful to the Genotoul bioinformatics platform Toulouse Occitanie (Bioinfo Genotoul, https://doi.org/10.15454/1.5572369328961167E12) for providing help and/or computing and/or storage resource. BIOGER benefits from the support of Saclay Plant Sciences-SPS (ANR-17-EUR-0007).

We also thank the BARIC workgroup (https://www.cesgo.org/catibaric/) for providing storage and computational resources. Rothamsted Research (JR, DS and RK) co-authors were supported by the Biotechnology and Biological Scientific Research Council (BBSRC) of the United Kingdom through the institute strategic grants “20:20 Wheat” and “Designing Future Wheat” (grant numbers BB/J/00426X/1 and BBS/E/C000I0250). The work (proposal: 10.46936/10.25585/60008023) conducted by the U.S. Department of Energy Joint Genome Institute (https://ror.org/04xm1d337), a DOE Office of Science User Facility, is supported by the Office of Science of the U.S. Department of Energy operated under Contract No. DE-AC02-05CH11231.

