## Supplementary figures and tables for "Improved gene annotation of the fungal wheat pathogen *Zymoseptoria tritici* based on combined Iso-Seq and RNA-Seq evidence"

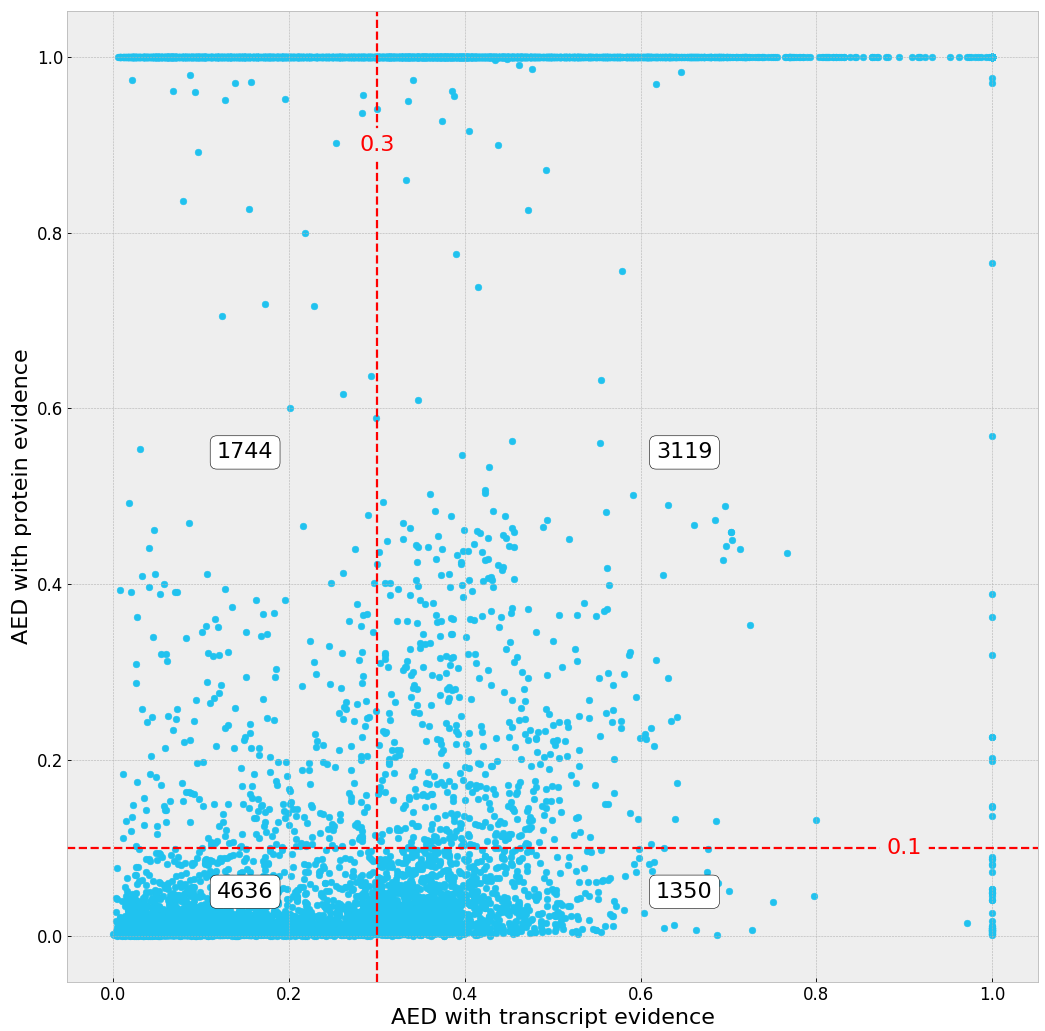

JGI

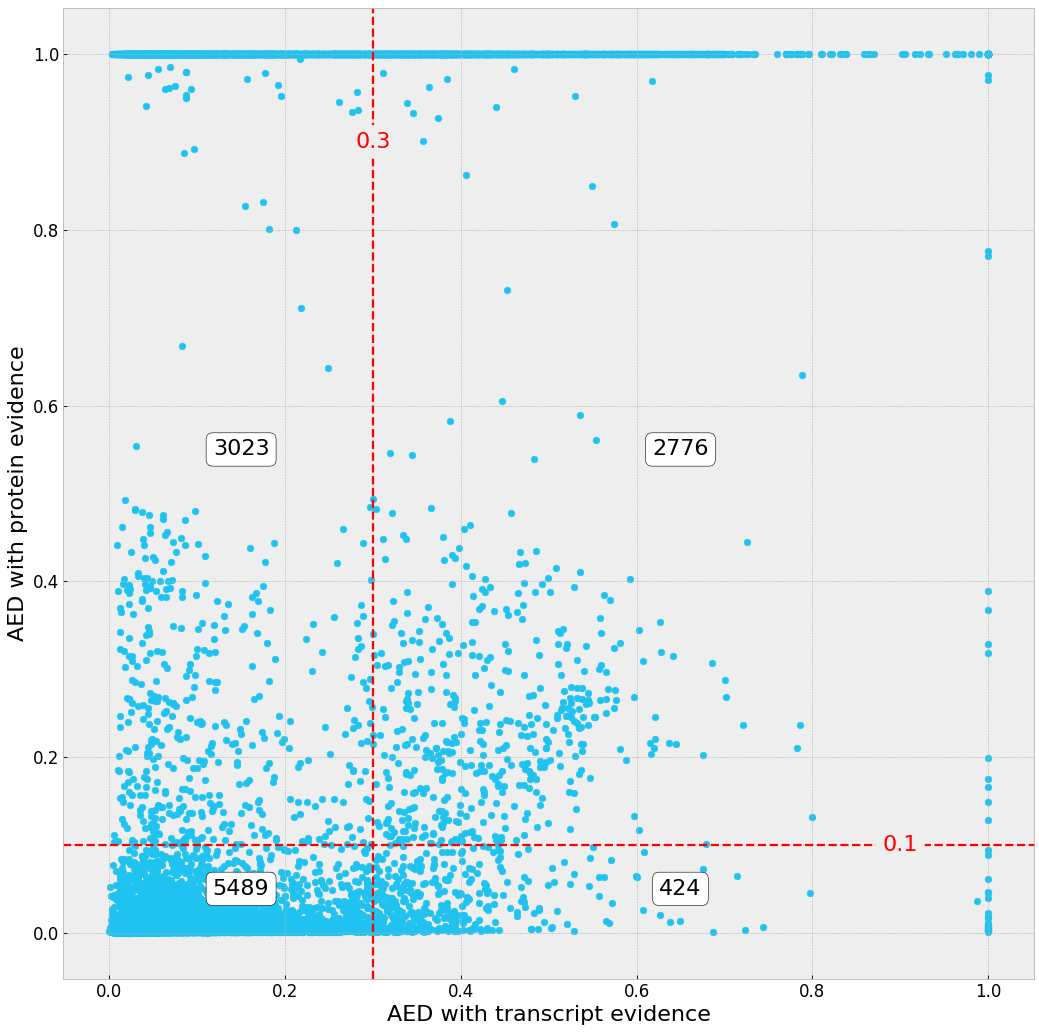

MPI

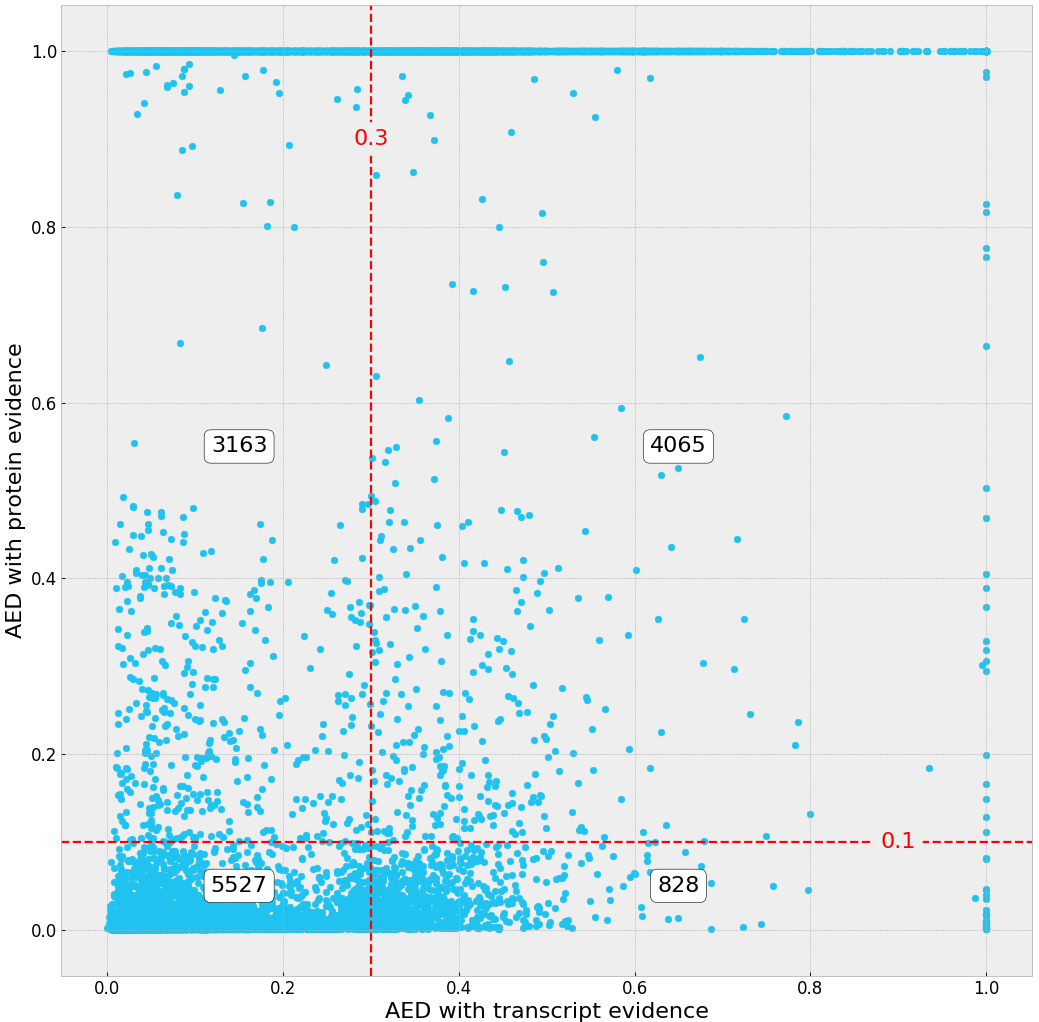

RRES

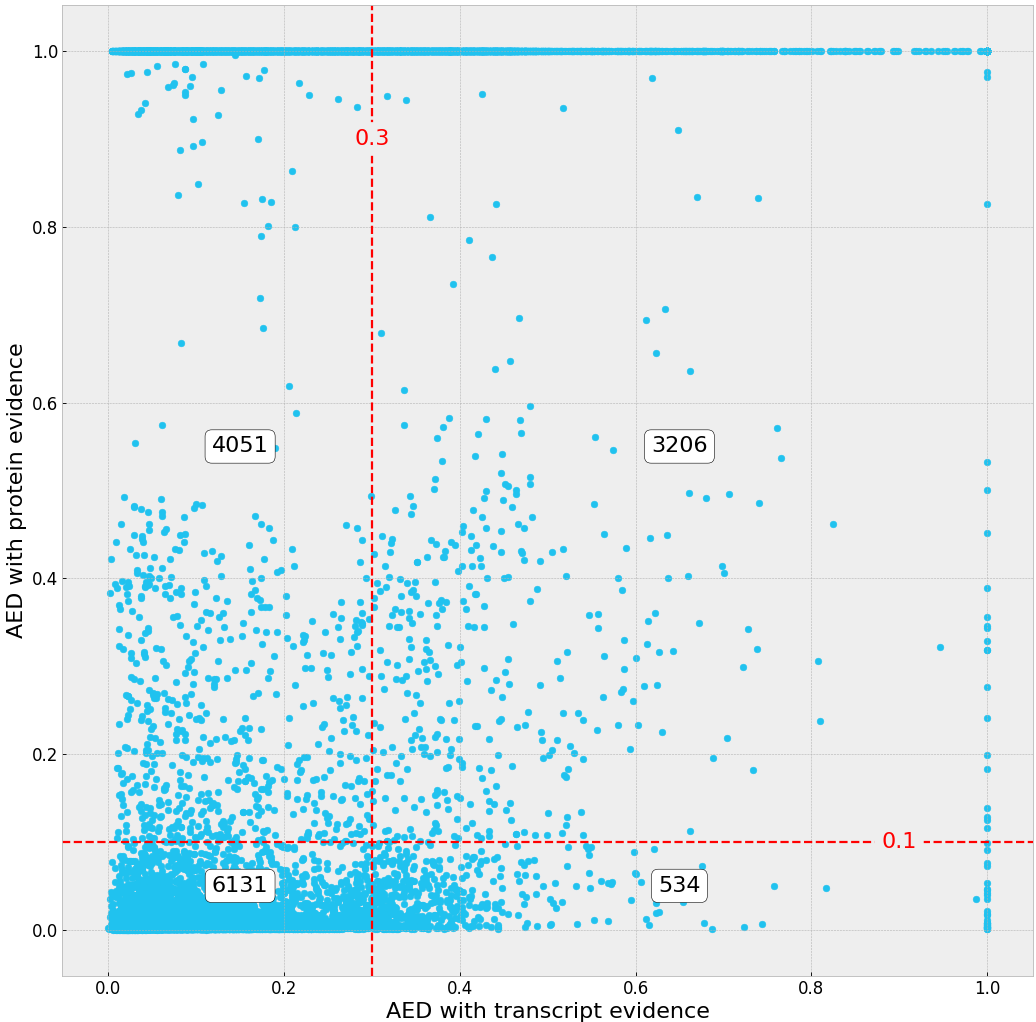

CURTIN

**Figure S1**. **Comparison of the Annotation Edit Distance (AED) scores of the JGI, MPI, RRES and CURTIN gene models**

AED scores (0-1) described how a given gene model fit the transcript and protein evidence (best fit = 0). Transcript evidence was computed from RNA-Seq and Iso-Seq data (X axis). Protein evidence was computed from fungal protein sequences excluding *Zymoseptoria* species (Y axis). The red, dashed lines represent the AED thresholds to filter out genes (0.3 for transcripts, 0.1 for proteins), except if they are supported by at least 4 different annotations (upper right area of the graph). The numbers of genes from each area were displayed in white boxes.

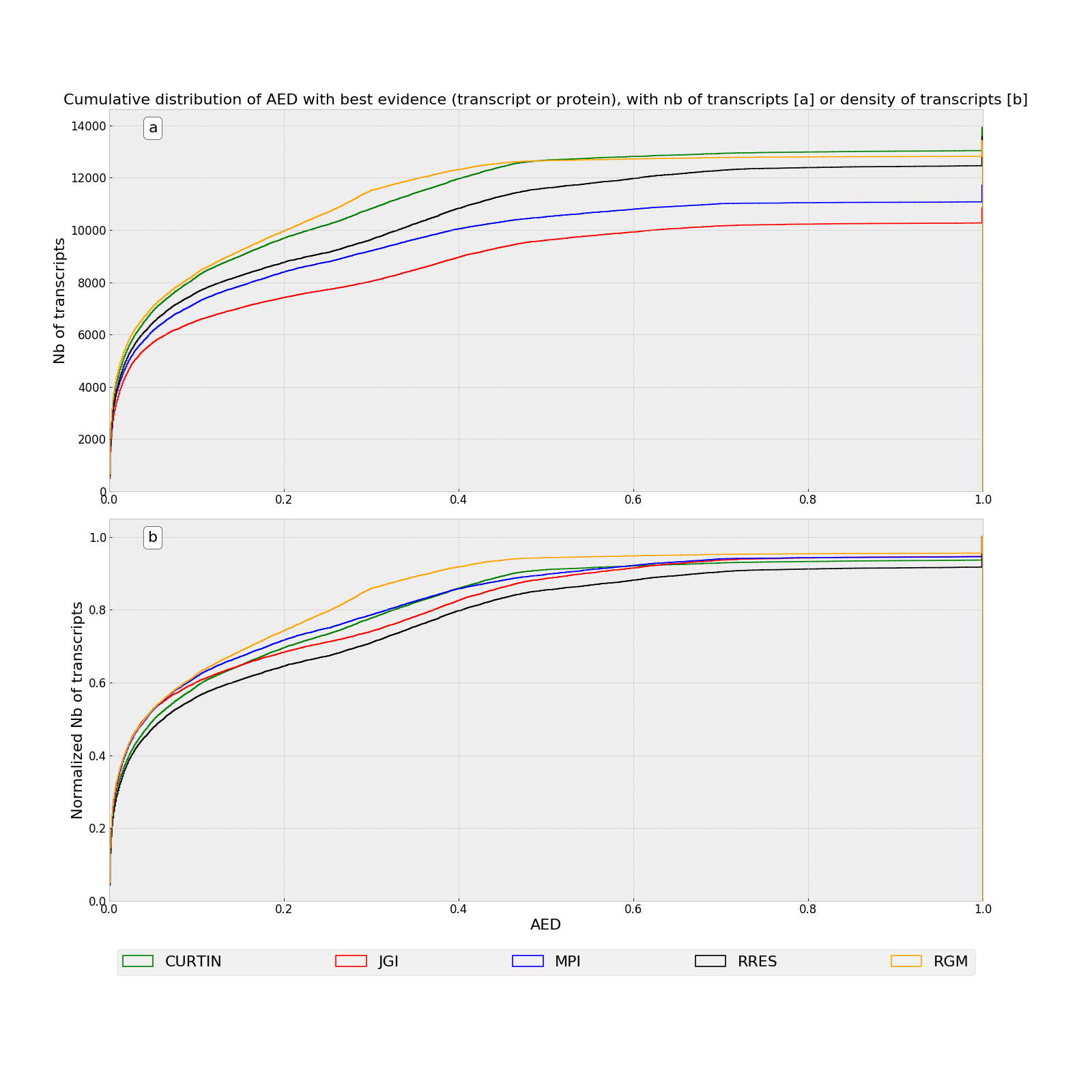

**Figure S2.** **Cumulative distributions of the best Annotation Edit Distance (AED) scores for Re-annotated Gene Models (RGMs) and those from previous annotations.**

AED scores (0-1) described how a given gene model fit the transcript and protein evidence (best fit = 0). The best AED score (X-axis) was computed from either transcript or protein evidence. a) cumulative plot of the number of transcripts, b) cumulative plot of the density of transcripts (normalized). The red line indicated the cutoff used to select the best gene model (0.3 for transcript evidence).

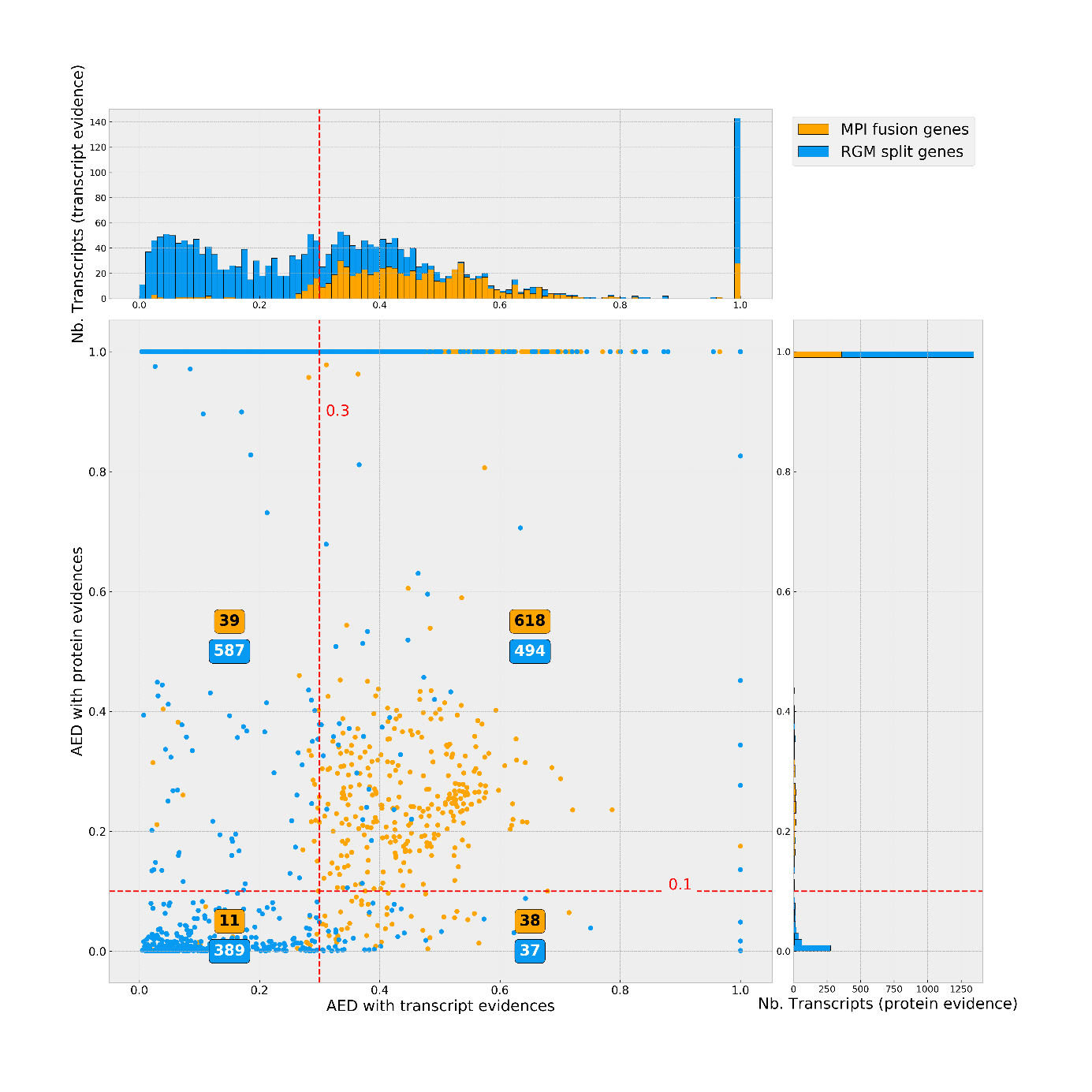

**Figure S3.** **Comparison of the Annotation Edit Distance (AED) scores of split Re-annotated Gene Models (RGMs) and their corresponding fused gene models from the MPI annotation**

AED scores (0-1) described how a given gene model fit the transcript and protein evidence (best fit = 0). Split RGMs were displayed in blue and the corresponding MPI fused genes were displayed in orange. Transcript evidence were computed from RNA-Seq or Iso-Seq data (X axis). Protein evidence were computed from fungal protein sequences excluding *Zymoseptoria* species (Y axis). The red, dashed lines corresponded to the AED thresholds used to filter out gene models (0.3 for transcripts, 0.1 for proteins). The number of transcripts for each value of AED score was plotted on cumulative histograms above the scatter plot (RGM in blue, MPI in orange). The number of transcripts with protein evidence were plotted on cumulative histograms on the right of the scatter plot (RGM in blue, MPI in orange).

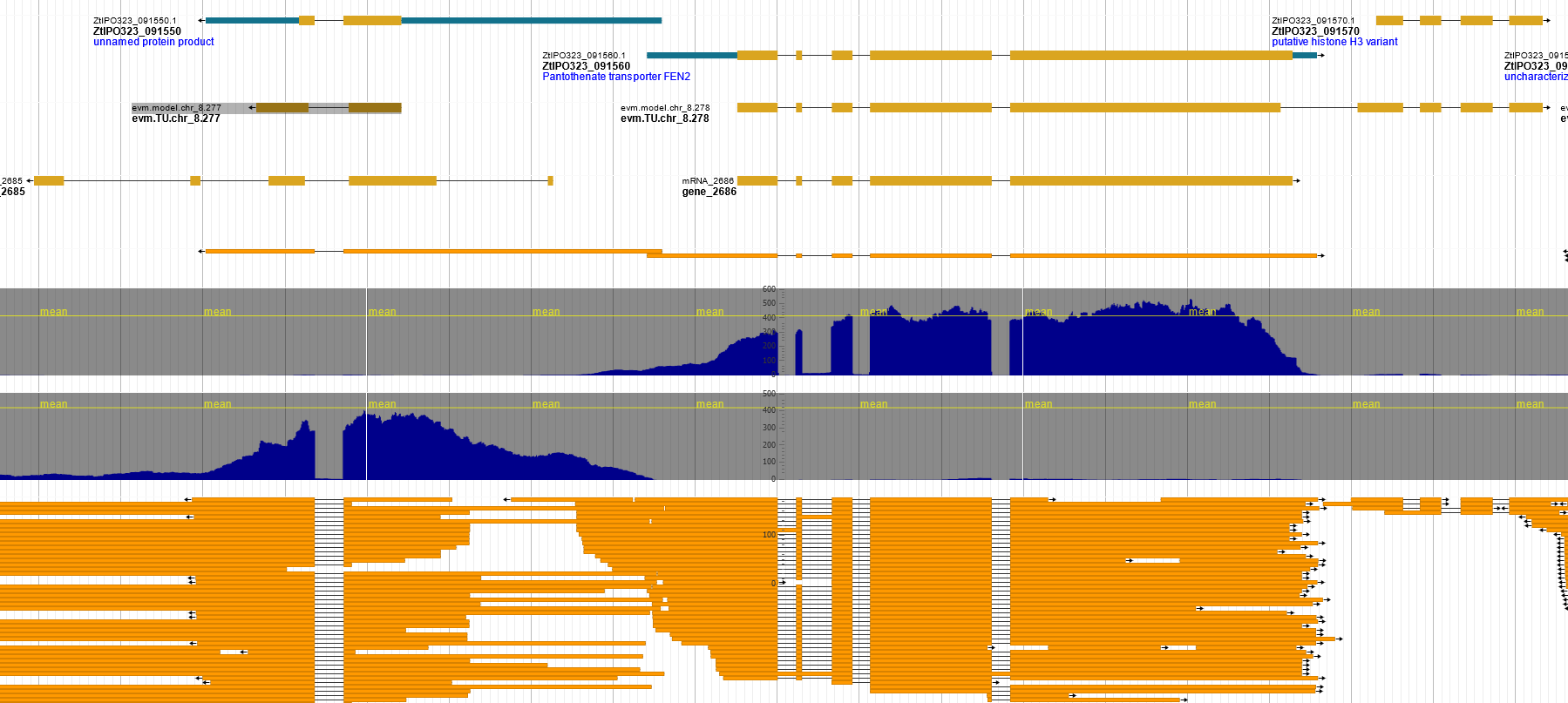

RGM

MPI

JGI

Iso-Seq

RNA-Seq

forward

RNA-Seq reverse

RNA-Seq transcripts

RGM-3

RGM-2

RGM-1

**Figure S4.** **Fused genes from the MPI annotation located at chr_8:940236...948036 splitted in two Re-annotated Gene Models (RGM-1 and RGM-2).**

In the region of chr_8:940236...948036 (7.8 Kb), three RGMs were predicted (red-dashed squares: RGM-1, RGM-2 and RGM-3). The MPI annotation predicted a gene model corresponding to the fusion of RGM2 and RGM-3 (green rectangle). This fusion had no transcript evidence (Iso-seq, RNA-Seq). Specific transcripts of RGM-2 (Iso-Seq, RNAseq) and RGM-3 (RNAseq) were detected. On the other strand, RGM-1 had correctly predicted intron splicing sites, while the corresponding gene models from the MPI and JGI annotations had incorrect intron splicing sites. RGM-3 was not predicted by JGI, despite encoding a conserved Histone-3 variant protein. RGM track: RGM gene models. MPI track: MPI gene models. JGI track: JGI gene models. Iso-Seq track: filtered Iso-Seq transcripts. RNA-Seq forward track: coverage of forward-strand RNA-Seq reads. RNA-Seq reverse track: coverage of reverse-strand RNA-Seq reads mapping at this locus. RNA-Seq transcript track: assembled RNA-Seq transcripts.

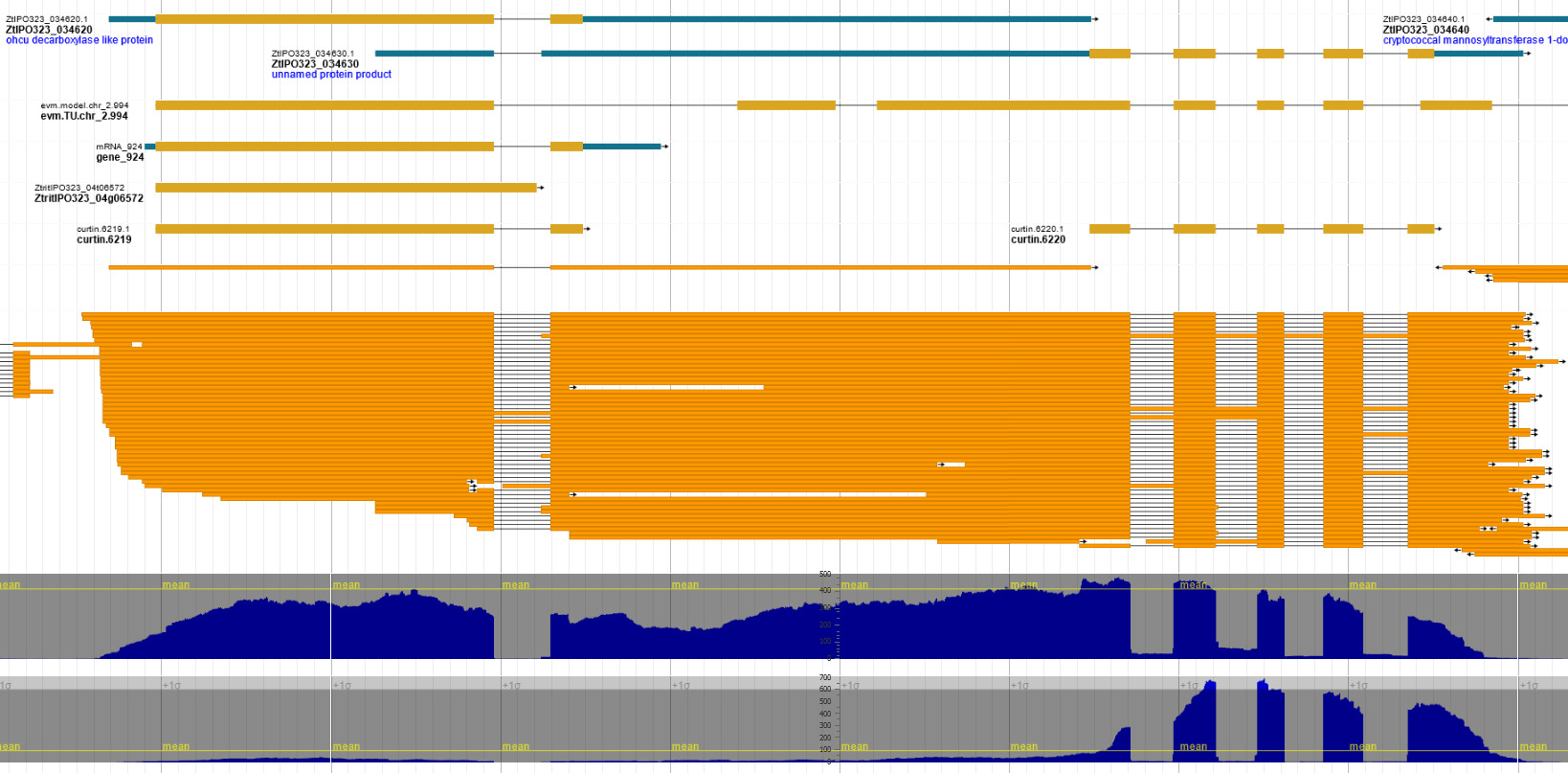

RGM

MPI

JGI

RRES

CURTIN

Iso-Seq

RNA-Seq, transcripts

RNA-Seq

coverage

Xylose

RNA-Seq

coverage

infection

13 dpi

RGM-1

RGM-2

**Figure S5.** **Fused genes from the MPI annotation located at chr_2:2938368...2940768 splitted in two Re-annotated Gene Models (RGM-1 and RGM-2)**

In the region of chr_2:2938368...2940768 (2.4 Kb), two RGMs were predicted (red-dashed squares). The MPI annotation predicted a gene model corresponding to the fusion of RGM-1 and RGM-2 (green rectangle) not supported by Iso-Seq. Iso-Seq transcripts supporting RGM-1 were detected. RGM-2 with no Iso-seq support was not predicted by JGI and RRES, while it was predicted as an independent gene in Curtin annotation. ). RGM2 (ZtIPO323_034630) encoded a Small Secreted Protein (SSP, see File S1). Assembled RNA-seq transcripts corresponding to the fused MPI gene model were detected. They were likely artefacts from assembly of overlapping RNA-Seq reads. Indeed, in a specific condition (infection at 13 days post inoculation), only RGM-2 transcript was detected (RNA-seq coverage 13 dpi track), supporting the prediction of RGM-2. RGM track: RGM gene models. MPI track: MPI gene model. JGI track: JGI gene model. RRES track: RRES gene model. Curtin track: Curtin gene models. Iso-Seq track: filtered Iso-Seq transcripts. RNA-Seq transcript track: assembled RNA-Seq transcripts. RNA-seq coverage Xylose: coverage of strand-specific RNA-Seq reads from a Xylose medium library. RNA-seq coverage infection 13 dpi: coverage of strand-specific RNA-Seq reads from a 13 dpi wheat infection library.

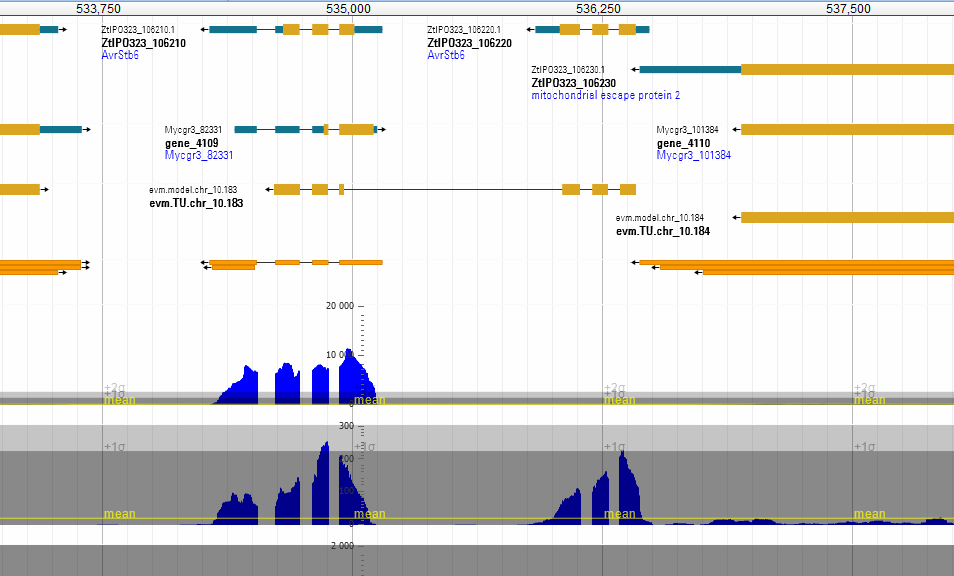

RNA-Seq

coverage

Glucose

Iso-Seq

MPI

JGI

RGM

RNA-Seq

coverage

infection 11 dpi

**Figure S6a. *Avr-Stb6* paralogs located on chromosome 10 predicted by the new annotation.**

The two new paralogs of *Avr-Stb6* (ZtIPO323_106210, ZtIPO323_106220) are located head to tail on chromosome 10 between position 534287 and 536486. ZtIPO323_106210 was predicted in JGI and MPI annotation, but the gene models did not match Iso-Seq and RNA-seq evidence. ZtIPO323_106220 was not predicted by any previous annotations. RGM annotation was only supported by RNA-seq evidence. ZtIPO323_106210 is expressed during *in vitro* growth ( RNA-Seq coverage glucose track), and infection stages (RNA-Seq coverage infection 11 dpi track). ZtIPO323_106220 was differentially upregulated during infection (RNA-Seq coverage infection 11 dpi track) to the same level as ZtIPO323_106210. RGM track: RGM gene models. JGI track: JGI gene model. MPI track: MPI gene model. Iso-Seq track: filtered Iso-Seq transcripts. RNA-seq coverage Glucose: coverage of strand-specific RNA-Seq reads from a Glucose medium library. RNA-seq coverage infection 11 dpi: coverage of strand-specific RNA-Seq reads from a 11 dpi wheat infection library.

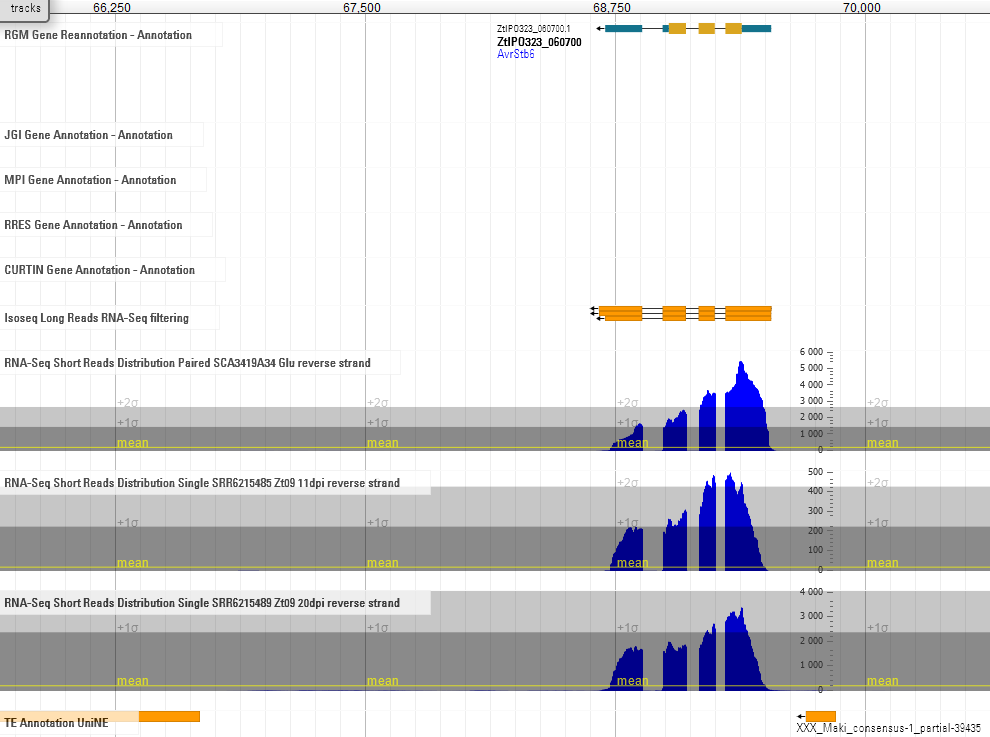

**Figure S6b. Original *Avr-Stb6* located on chromosome 5**

The original *Avr-Stb6* is located at the end of chromosome 5. It was not predicted by any previous annotations, while it was correctly predicted as RGM (see track RGM track reannotation). Iso-Seq transcripts were detected (see track Isoseq long reads), since this gene is highly expressed during *in vitro* growth of *Zymoseptoria tritici*, as well as during infection (see tracks RNA-seq short reads distribution). RGM track: RGM gene models. JGI track: JGI gene model. MPI track: MPI gene model. RRES track: RRES gene model. Curtin track: Curtin gene models. Iso-Seq track: filtered Iso-Seq transcripts. RNA-seq distribution Glucose: coverage of strand-specific RNA-Seq reads from a Glucose medium library. RNA-seq distribution infection 11 dpi: coverage of strand-specific RNA-Seq reads from a 11 dpi wheat infection library.

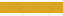

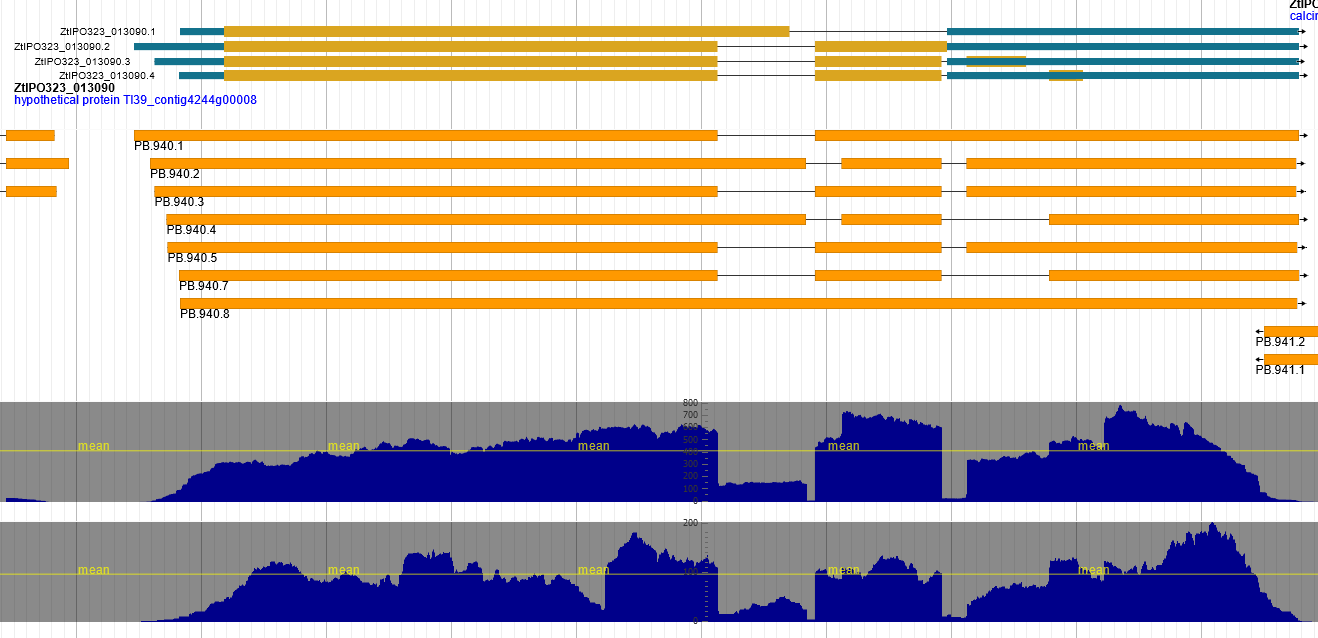

RGM

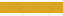

Iso-Seq

RNA-Seq

coverage

Xylose

RNA-Seq

coverage

AE

**Figure S7. Isoforms of Re-annotated Gene Model (RGM) ZtIPO323_013090 supported by Iso-Seq**

RGM ZtIPO323_013090 located at chr_1:3358097...3361350 (3.25 Kb) has four different isoforms. One alternative splicing site (red-dashed rectangle) supported by RNA-Seq, was not supported by Iso-Seq. Another alternative splicing site detected both by RNA-Seq and Iso-Seq (black-dashed rectangle), was not used to predict an isoform by ab initio software due to a stop codon before the splicing site. Finally, the canonical form retained is the transcript without any introns. This selection could be an artefact of transcripts from AE medium inducing many intron retention events. The canonical isoform is likely the RGM corresponding to the Iso-Seq transcript PB.940.5 with two introns (isoform 3). RGM track: RGM gene model. Iso-Seq track: filtered Iso-Seq transcripts. RNA-Seq coverage Xyl track: coverage of strand-specific RNA-Seq reads from the Xylose medium library. RNA-Seq coverage AE track: coverage of strand-specific RNA-Seq reads from AE medium library.

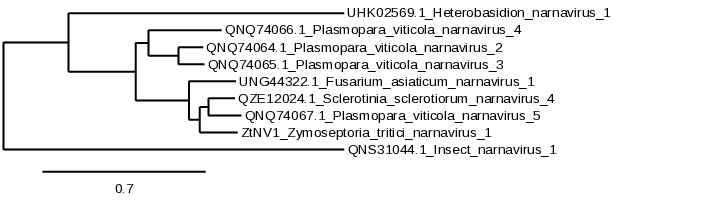

**100**

**85**

**93**

**100**

**Figure S8.** **Phylogenetic tree of RNA-dependent RNA polymerases of fungal narnaviruses related Zt-NV1 from *Zymoseptoria tritici***

RNA-dependent RNA polymerase sequences from narnaviruses related to Zt-NV1 (blue rectangle) were retrieved from NCBI. Phylogenetic analysis was performed with PhyML 3.1 at Phylo.fr (Dereeper et al., 2008). Bootstrap values over 50% are indicated on supported branches (1000 replicates).

| Category | JGI | MPI | RRES | CURTIN |
| --- | --- | --- | --- | --- |
| nb_CDS | 10849 | 11712 | 13583 | 13922 |
| average_CDS_length, bp | 1307 | 1465 | 1293 | 1287 |
| median_CDS_length, bp | 1071 | 1203 | 1044 | 1041 |
| min_CDS_length, bp | 150 | 150 | 96 | 93 |
| max_CDS_length, bp | 13842 | 18297 | 18423 | 14523 |
| nb_exons | 28313 | 29728 | 30772 | 30564 |
| average_exons_per_CDS | 2.6 | 2.5 | 2.2 | 2.2 |
| average_exon_length, bp | 531 | 577 | 570 | 586 |
| min_exon_length | 2 | 1 | 1 | 1 |
| max_exon_length | 12888 | 12975 | 18423 | 9987 |
| nb_transcript_mono_exon | 3153 | 3746 | 5233 | 5594 |
| nb_introns | 17464 | 18016 | 17189 | 16642 |
| average_introns_per_transcript | 1.6 | 1.5 | 1.2 | 1.2 |
| average_intron_length | 133 | 93 | 109 | 92 |
| min_intron_length | 11 | 23 | 4 | 10 |
| max_intron_length | 42135 | 7292 | 59574 | 5000 |

**Table S1.** **CDS features of the four available gene annotations of the *Z. tritici* IPO323 genome (JGI, MPI, RRES and CURTIN).**

Gene models were filtered out for transposable elements. nb : number, min : minimum, max : maximum.

|  | JGI | | MPI | | RRES | | CURTIN | |
| --- | --- | --- | --- | --- | --- | --- | --- | --- |
| Chr | #genes | % | #genes | % | #genes | % | #genes | % |
| 1 | 1975 | 18,2% | 2107 | 18,0% | 2321 | 17,1% | 2326 | 17,8% |
| 2 | 1127 | 10,4% | 1236 | 10,6% | 1377 | 10,1% | 1380 | 10,5% |
| 3 | 1067 | 9,8% | 1138 | 9,7% | 1297 | 9,5% | 1261 | 9,6% |
| 4 | 818 | 7,5% | 889 | 7,6% | 998 | 7,3% | 993 | 7,6% |
| 5 | 776 | 7,2% | 848 | 7,2% | 986 | 7,3% | 971 | 7,4% |
| 6 | 685 | 6,3% | 716 | 6,1% | 820 | 6,0% | 810 | 6,2% |
| 7 | 758 | 7,0% | 842 | 7,2% | 975 | 7,2% | 957 | 7,3% |
| 8 | 685 | 6,3% | 749 | 6,4% | 843 | 6,2% | 817 | 6,2% |
| 9 | 601 | 5,5% | 620 | 5,3% | 703 | 5,2% | 689 | 5,3% |
| 10 | 513 | 4,7% | 551 | 4,7% | 624 | 4,6% | 616 | 4,7% |
| 11 | 482 | 4,4% | 523 | 4,5% | 604 | 4,4% | 607 | 4,6% |
| 12 | 405 | 3,7% | 446 | 3,8% | 514 | 3,8% | 523 | 4,0% |
| 13 | 325 | 3,0% | 362 | 3,1% | 411 | 3,0% | 401 | 3,1% |
| 14 | 111 | 1,0% | 108 | 0,9% | 163 | 1,2% | 115 | 0,9% |
| 15 | 83 | 0,8% | 84 | 0,7% | 143 | 1,1% | 99 | 0,8% |
| 16 | 86 | 0,8% | 102 | 0,9% | 169 | 1,2% | 98 | 0,7% |
| 17 | 76 | 0,7% | 84 | 0,7% | 131 | 1,0% | 74 | 0,6% |
| 18 | 60 | 0,6% | 75 | 0,6% | 121 | 0,9% | 91 | 0,7% |
| 19 | 84 | 0,8% | 78 | 0,7% | 141 | 1,0% | 90 | 0,7% |
| 20 | 78 | 0,7% | 89 | 0,8% | 135 | 1,0% | 92 | 0,7% |
| 21 | 54 | 0,5% | 65 | 0,6% | 107 | 0,8% | 84 | 0,6% |
| Total | 10849 |  | 11712 |  | 13583 |  | 13094 |  |

**Table S2.** **Chromosome localization of gene models of the four available annotations of the *Z. tritici* IPO323 genome (JGI, MPI, RRES and CURTIN).**

Chr: Chromosome, of which the first 13 are the core and the remaining 8 accessories.

#genes: ratio compared to the whole dataset.

| Condition | Replicate | Type | # transcripts -post stringtie | # transcripts - post jaccardclip | % clipped reads | # transcripts - post cov filter | RUN accession | Corresponding Isoseq accession |
| --- | --- | --- | --- | --- | --- | --- | --- | --- |
| AE (Yeast extract Glycerol) | 1 | PE-stranded | 11686 | 12528 | 4,40% | 12369 | GSM6758342 | GSM6758369 |
| AE (Yeast extract Glycerol) | 2 | PE-stranded | 10937 | 11803 | 5,10% | 11644 | GSM6758343 |  |
| AE (Yeast extract Glycerol) | 3 | PE-stranded | 10967 | 11666 | 4,30% | 11544 | GSM6758344 |  |
| MMZt NO3 + Glucose | 1 | PE-stranded | 10780 | 11568 | 5,10% | 11406 | GSM6758345 | GSM6758370 |
| MMZt NO3 + Glucose | 2 | PE-stranded | 9713 | 10432 | 5,10% | 10303 | GSM6758346 |  |
| MMZt NO3 + Glucose | 3 | PE-stranded | 10807 | 11796 | 5,90% | 11595 | GSM6758347 |  |
| MMZt NO3 + Sucrose | 1 | PE-stranded | 10796 | 11683 | 5,60% | 11522 | GSM6758348 | NA |
| MMZt NO3 + Sucrose | 2 | PE-stranded | 10876 | 11648 | 4,70% | 11511 | GSM6758349 |  |
| MMZt NO3 + Sucrose | 3 | PE-stranded | 10496 | 11428 | 5,60% | 11301 | GSM6758350 |  |
| MMZt NO3 + Xylose | 1 | PE-stranded | 10881 | 11913 | 6,20% | 11722 | GSM6758351 | GSM6758371 |
| MMZt NO3 + Xylose | 2 | PE-stranded | 10575 | 11296 | 4,70% | 11135 | GSM6758352 |  |
| MMZt NO3 + Xylose | 3 | PE-stranded | 10636 | 11310 | 4,30% | 11165 | GSM6758353 |  |
| MMZt NO3 + Mannitol | 1 | PE-stranded | 10534 | 11275 | 4,60% | 11142 | GSM6758354 | GSM6758372 |
| MMZt NO3 + Mannitol | 2 | PE-stranded | 11008 | 12038 | 6,00% | 11832 | GSM6758355 |  |
| MMZt NO3 + Mannitol | 3 | PE-stranded | 10848 | 11693 | 5,20% | 11539 | GSM6758356 |  |
| MMZt NO3 + Galactose | 1 | PE-stranded | 10831 | 12045 | 7,10% | 11831 | GSM6758357 | NA |
| MMZt NO3 + Galactose | 2 | PE-stranded | 10939 | 11950 | 6,00% | 11794 | GSM6758358 |  |
| MMZt NO3 + Galactose | 3 | PE-stranded | 11000 | 12064 | 6,30% | 11853 | GSM6758359 |  |
| MMZt NO3_Glucose +SAHA | 1 | PE-stranded | 11141 | 12344 | 7,00% | 12132 | GSM6758360 | GSM6758373 |
| MMZt NO3_Glucose +SAHA | 2 | PE-stranded | 11199 | 12422 | 6,80% | 12226 | GSM6758361 |  |
| MMZt NO3_Glucose +SAHA | 3 | PE-stranded | 11283 | 13039 | 8,90% | 12767 | GSM6758362 |  |
| MMZt NO3_Glucose +TSA | 1 | PE-stranded | 11473 | 13061 | 8,30% | 12785 | GSM6758363 | GSM6758374 |
| MMZt NO3_Glucose +TSA | 2 | PE-stranded | 11762 | 13325 | 8,10% | 13048 | GSM6758364 |  |
| MMZt NO3_Glucose +TSA | 3 | PE-stranded | 11566 | 13287 | 8,80% | 13021 | GSM6758365 |  |
| YPD - 18 °C | 1 | PE-stranded | 10964 | 12468 | 8,20% | 12240 | GSM6758366 | GSM6758378 |
| YPD - 18 °C | 2 | PE-stranded | 11018 | 12442 | 7,70% | 12201 | GSM6758367 |  |
| YPD - 18 °C | 3 | PE-stranded | 10923 | 12462 | 8,30% | 12228 | GSM6758368 |  |
| Wheat infection - 4 dpi | 1 | SE-stranded | 10349 | 10431 | 0,77% | 10395 | SRR6215483 |  |
| Wheat infection - 4 dpi | 2 | SE-stranded | 8661 | 8671 | 0,12% | 8665 | SRR6215484 |  |
| Wheat infection - 11 dpi | 1 | SE-stranded | 9135 | 9141 | 0,07% | 9133 | SRR6215485 |  |
| Wheat infection - 11 dpi | 2 | SE-stranded | 9803 | 9816 | 0,13% | 9804 | SRR6215486 |  |
| Wheat infection - 13 dpi | 1 | SE-stranded | 11675 | 11688 | 0,09% | 11672 | SRR6215487 |  |
| Wheat infection - 13 dpi | 2 | SE-stranded | 11357 | 11362 | 0,04% | 11348 | SRR6215488 |  |
| Wheat infection - 20 dpi | 1 | SE-stranded | 12215 | 12226 | 0,07% | 12205 | SRR6215489 |  |
| Wheat infection - 20 dpi | 2 | SE-stranded | 12196 | 12202 | 0,04% | 12180 | SRR6215490 |  |
| YMS - 18 °C | 1 | SE-stranded | 9183 | 11100 | 11,80% | 10952 | SRR8788920 |  |
| YMS - 18 °C | 2 | SE-stranded | 8885 | 10930 | 13,10% | 10797 | SRR8788921 |  |
| YMS - 18 °C kmt1 mutant | 1 | SE-stranded | 9696 | 12054 | 13,00% | 11875 | SRR8788922 |  |
| YMS - 18 °C kmt1 mutant | 2 | SE-stranded | 9554 | 11725 | 12,20% | 11559 | SRR8788923 |  |
| YMS - 18°C kmt6 mutant | 1 | SE-stranded | 8688 | 10876 | 13,30% | 10720 | SRR8788924 |  |
| YMS - 18°C kmt6 mutant | 2 | SE-stranded | 9001 | 11152 | 13,20% | 11006 | SRR8788925 |  |
| YMS - 18 °C kmt1/6 mutant | 1 | SE-stranded | 10408 | 13168 | 13,30% | 12929 | SRR8788926 |  |
| YMS - 18 °C kmt1/6 mutant | 2 | SE-stranded | 10415 | 13127 | 13,10% | 12914 | SRR8788927 |  |
| YPD-25°C |  |  |  |  |  |  |  | GSM6758379 |
| PDB-18°C |  |  |  |  |  |  |  | GSM6758375 |
| PDB-25°C |  |  |  |  |  |  |  | GSM6758376 |

**Table S3.** **RNA-Seq and Iso-Seq cDNA libraries from *Z. tritici* IPO323**

Ten growth conditions were used to produce RNAs for Iso-seq and RNA-seq libraries. Additional single-stranded RNA-seq data were obtained from public databases (see below). RNA-Seq sequences were used for transcript assemblies and differential expression analyses. Iso-Seq sequences were used for gene annotation and isoform analyses. Wheat infection data for SRR6215483- SRR6215490 and YMS mutants SRR8788920- SRR8788927 were downloaded from the Sequence Read Archive (SRA) repository. Other RNA-Seq data generated in this study were submitted to the SRA database (see columns accession).

| ## statistics ## |  |
| --- | --- |
| nb_genes | 13414 |
| average_gene_length | 1900 |
| median_gene_length | 1635 |
| min_gene_length | 102 |
| max_gene_length | 17065 |
| nb_transcripts | 13414 |
| average_transcripts_per_gene | 1 |
| average_transcript_length | 1900 |
| median_transcript_length | 1635 |
| min_transcript_length | 102 |
| max_transcript_length | 17065 |
| nb_exons | 30946 |
| average_exons_per_transcript | 2.3 |
| average_exon_length | 782 |
| median_exon_length | 431 |
| min_exon_length | 1 |
| max_exon_length | 16680 |
| nb_transcript_mono_exon | 4850 |
| nb_introns | 17532 |
| average_introns_per_transcript | 1.3 |
| average_intron_length | 73 |
| median_intron_length | 57 |
| min_intron_length | 5 |
| max_intron_length | 3166 |
| nb_CDS | 13414 |
| average_CDS_length | 1287 |
| median_CDS_length | 1041 |
| min_CDS_length | 102 |
| max_CDS_length | 16506 |
| nb_transcripts_with_utr | 9856 |
| average_five_prime_utr_length | 315 |
| median_five_prime_utr_length | 156 |
| min_five_prime_utr_length | 1 |
| max_five_prime_utr_length | 7053 |
| average_three_prime_utr_length | 389 |
| median_three_prime_utr_length | 220 |
| min_three_prime_utr_length | 1 |
| max_three_prime_utr_length | 8647 |

**Table S4. Features of Re-annotated Gene Models (RGMs) of *Z. tritici* IPO323 genome**

nb : number, min : minimum, max : maximum, utr : untranslated region from transcripts

| BUSCO category | JGI | MPI | CURTIN | RRES | RGM |
| --- | --- | --- | --- | --- | --- |
| Complete BUSCOs (C) | 1633 | 1679 | 1681 | 1693 | 1696 |
| Complete BUSCOs (C) % | 95.7% | 98.4% | 98.5% | 99.2% | 99.4% |
| Complete and single-copy BUSCOs (S) | 1632 | 1678 | 1615 | 1692 | 1695 |
| Complete and duplicated BUSCOs (D) | 1 | 1 | 66 | 1 | 1 |
| Fragmented BUSCOs (F) | 25 | 3 | 8 | 5 | 2 |
| Missing BUSCOs (M) | 48 | 24 | 17 | 8 | 8 |
| Total BUSCO groups | 1706 | | | | |

**Table S5.** **Benchmarking Universal Single-Copy Orthologs (BUSCO) analysis of Re-annotated Gene Models (RGM) and gene models from previous annotations of the *Z. tritici* IPO323 genome.**

This comparison was performed using BUSCO and the ascomycota_odb as reference genes. Higher BUSCO scores (99.4 % identical) were obtained with RGMs compared to the JGI, MPI and CURTIN annotations (95.7-98.5% identical), while scores obtained with RRES gene models were similar to RGMs (99.1 % identical). The JGI annotation had the highest number of missing BUSCO genes.

| Chromosome | # RGMs | # dissimilar RGMs | # specific RGMs | # RGMs with no evidence |
| --- | --- | --- | --- | --- |
| 1 | 2365 | 184 | 100 | 47 |
| 2 | 1392 | 132 | 54 | 28 |
| 3 | 1291 | 130 | 61 | 32 |
| 4 | 1021 | 70 | 54 | 29 |
| 5 | 997 | 98 | 49 | 18 |
| 6 | 835 | 78 | 41 | 12 |
| 7 | 952 | 88 | 39 | 224 |
| 8 | 843 | 79 | 33 | 27 |
| 9 | 708 | 68 | 34 | 25 |
| 10 | 624 | 56 | 21 | 29 |
| 11 | 604 | 53 | 28 | 9 |
| 12 | 542 | 51 | 39 | 11 |
| 13 | 423 | 42 | 23 | 12 |
| 14 | 133 | 41 | 18 | 8 |
| 15 | 115 | 43 | 18 | 7 |
| 16 | 103 | 28 | 10 | 15 |
| 17 | 89 | 24 | 9 | 8 |
| 18 | 94 | 26 | 5 | 8 |
| 19 | 103 | 30 | 19 | 9 |
| 20 | 93 | 23 | 7 | 8 |
| 21 | 87 | 32 | 9 | 8 |
| Total | 13414 | 1376 | 671 | 574 |

**Table S6.** **Distribution of Re-annotated Gene Models (RGMs) on *Z. tritici* IPO323 chromosomes**

\# RGMs: number of Re-annotated Gene Models (RGMs).

\# dissimilar RGMs: number of RGMs with the following properties, at a given locus, a RGM was predicted by at least one other annotation, but they differed in their structure, here RGMs differing from all previous annotations.

\# specific RGMs: number of RGMs with the following properties, at a given locus, a single RGM is predicted by a single annotation, here RGM specific.

\# RGMs with no evidence: number of RGMs with the following properties, RGMs without transcript or protein evidence, but rescued since they were predicted by at least four different annotations.

| fusion / split | JGI | MPI | CURTIN | RRES | RGM |
| --- | --- | --- | --- | --- | --- |
| JGI | — | 312/674 | 626/1385 | 425/946 | **558/1258** |
| MPI | 286/584 | — | 801/1702 | 533/1127 | **706/1507** |
| CURTIN | 113/230 | 31/63 | — | 133/278 | **83/176** |
| RRES | 102/206 | 51/103 | 445/929 | — | **333/701** |
| RGM | **92/186** | **19/38** | **177/363** | **98/200** | — |

**Table S7. Identification of fused/split genes in the *Z. tritici* IPO323 genome annotations.**

These comparisons were performed in a pairwise manner. The first value is the number of fused genes for the horizontal entry and the second is the number of split genes for the vertical entry.

| Isoforms per RGM | 1 | 2 | 3 | 4 | 5 | 6+ |
| --- | --- | --- | --- | --- | --- | --- |
| Number of RGMs | 11672 | 1342 | 274 | 77 | 29 | 20 |

**Table S8.** **Number of transcript isoforms detected for Re-annotated Gene Models (RGMs) in *Z. tritici* IPO323**

The gene with the highest number of isoforms detected by Iso-Seq is ZtIPO323_108820 (chr_10:1162483...1166667 - 4.19 Kb) with 15 isoforms, among which only a few were supported quantitatively by RNA-Seq.

| LncRNA | Type | Up in planta log2FC > 2 | Down in planta log2FC < 2 | Gene (antisense) | Gene annotation |
| --- | --- | --- | --- | --- | --- |
| PB.854.1 | intergenic |  | Y |  |  |
| PB.927.1 | intergenic |  | Y |  |  |
| PB.1188.1 | antisens |  | Y | ZtIPO323_016330 | subtilisin-like protein |
| PB.1594.1 | antisens |  | Y | ZtIPO323_022040 | related to allantoase permease |
| PB.2214.1 | antisens |  | Y | ZtIPO323_030630/ ZtIPO323_030640 | NA |
| PB.2569.1 | antisens |  | Y | ZtIPO323_035460 | alpha/beta like hydrolase |
| PB.2709.1 | antisens | Y |  | ZtIPO323_037670 | TTL-domain containing protein |
| PB.4776.1 | antisens |  | Y | ZtIPO323_066740 | NA |
| PB.5130.1 | intergenic | Y |  |  |  |
| PB.5328.2 | antisens |  | Y | ZtIPO323_074720 | NA |
| PB.5366.1 | antisens |  | Y | ZtIPO323_075280 | HSP20-like chaperone |
| PB.6002.1 | antisens |  | Y | ZtIPO323_084180 | P450 monooxygenbase |
| PB.6788.1 | antisens | Y |  | UTR overlap |  |
| PB.7120.1 | antisens |  | Y | ZtIPO323_102870 | Phosphomevalonate kinase |
| PB.7618.1 | antisens |  | Y | ZtIPO323_110030 | glycoside hydrolase family 45 protein |
| PB.8769.1 | intergenic | Y |  |  |  |
| PB.8778.1 | intergenic | Y |  |  |  |

**Table S9. Long non-coding RNAs (lncRNA) from *Z. tritici* IPO323 differentially expressed during infection**

List of lncRNAs annotated from Iso-Seq data differentially expressed during infection with a strong fold change (4X). Antisense lncRNAs are preferentially involved in cis regulation, whereas intergenic lncRNAs are preferentially involved in trans regulation.

| Type | Annotation | Identical CDS* | | Unique identical CDS** | |
| --- | --- | --- | --- | --- | --- |
| Available annotations | JGI | 4865 | 11367 | 157 | 929 |
|  | MPI | 8431 |  | 91 |  |
|  | RRES | 8317 |  | 175 |  |
|  | CURTIN | 9584 |  | 506 |  |
| New annotations | Eugene | 10224 | 11677 | 1603 | 1802 |
|  | LoReAn | 7769 |  | 199 |  |

**Table S10.** **Contribution of each annotation of *Z. tritici* IPO323 genome to Re-annotated Gene Models (RGMs)**

The annotations that contributed the most to RGMs were respectively Eugene (76% Identical CDS*, 1603 Unique identical CDS**) and Curtin (71% Identical CDS, 506 Unique identical CDS). Combining gene models from the four available annotations (JGI, MPI, RRES, CURTIN) showed that 11367 of their CDS were identical to RGMs (contribution: 84.7%). Combining gene models from the two new annotations (Eugene, LoRean) showed that 11677 of their CDS were identical to RGMs (contribution: 87%). The combination of the six annotations was needed to predict all the 13414 RGMs.

* Identical coding sequence (CDS): number of CDS identical to RGMs

** Unique identical coding sequence (CDS): number of CDS predicted in a single annotation and retained as RGMs.
